# Mucin-enriched environment shapes bacteriophage protection against *Pseudomonas aeruginosa* in airway epithelial cells

**DOI:** 10.1101/2025.10.17.683030

**Authors:** Daniel de Oliveira Patricio, Katri Niitti, Lucilene Wildner Granella, Valtteri Suokko, Gabriel Magno de Freitas Almeida, Lotta-Riina Sundberg

**Affiliations:** Department of Biological and Environment Science, Nanoscience Center, University of Jyväskylä, Jyväskylä, Finland; The Norwegian College of Fishery Science, Faculty of Biosciences, Fisheries and Economics, UiT— The Arctic University of Norway, Tromsø, Norway

## Abstract

Mucin hypersecretion, a hallmark of chronic respiratory diseases (CRD), creates a complex microenvironment that reshapes host immunity and microbial behavior. However, its impact on bacteriophage therapy remains poorly understood. Here, we demonstrate that, despite reducing *Pseudomonas aeruginosa* internalization, exogenous mucin increases bacterial-induced cytotoxicity and inflammation in airway epithelial cells, while driving CRD-like transcriptional changes, including hypoxia and stress responses. Under bacterial culture, mucin selectively downregulates bacterial virulence factors without impairing growth. *P. aeruginosa*-infecting bacteriophage DMS3vir retains full lytic activity in mucin-rich conditions and, in synergy with mucin, enhances protection against cytotoxicity in epithelial cells. DMS3vir also reduces IL-8 gene expression without triggering antiviral responses. Furthermore, mucin supplementation shapes phage-resistant *P. aeruginosa* phenotypes, altering colony pigmentation, pyocyanin production, and motility, influencing virulence trade-offs. These findings uncover the dual role of mucins as modulators of infection and sensitizers to phage protection, paving the way for optimized, mucosa-adapted phage therapies in chronic lung diseases.

## 1. Introduction

Chronic respiratory diseases (CRD), including cystic fibrosis (CF), chronic obstructive pulmonary disease (COPD) and asthma, are characterized by persistent inflammation, impaired mucociliary clearance, and recurrent airway infections^1–3^. A hallmark of these conditions is the hypersecretion of mucins, gel-forming glycoproteins such as MUC5AC and MUC5B, which structure the mucosal layer, trap pathogens, and help maintain epithelial integrity^1–5^. Increased mucin production by goblet cells can be influenced by various mediators such as pathogens, inflammatory responses, and xenobiotic exposure^2^. Mucin overproduction alters the physical and biochemical properties of the airway surface, creating a viscous, hypoxic, and oxidatively stressed microenvironment that disrupt immune balance and facilitates microbial colonization^2,6–10^.

*Pseudomonas aeruginosa*, a key opportunistic pathogen found in chronic airway infections, thrives under hypoxic conditions. It contributes to epithelial injury and lung function decline through development of antimicrobial resistance, biofilm formation, and secretion of virulence factors regulated by quorum sensing (QS)^11,12^. The mucosal environment directly shapes *P. aeruginosa* behavior. For instance, mucin glycans have been shown to attenuate virulence by disrupting QS signaling and inhibiting biofilm formation^13,14^. Moreover, mucin hyper-concentration contributes to bacterial aggregation that increases antibiotic tolerance variability^15,16^. In addition to shaping microbial behavior, mucins are tightly linked to epithelial immune responses, particularly through the regulation of key pro-inflammatory cytokines^9,10^.

Pro-inflammatory cytokines such as interleukin-6 (IL-6) and IL-8 (or CXCL8) play central roles during chronic respiratory diseases and are associated with mucin hypersecretion^1,17^. IL-8 is induced by pathogen-associated molecular patterns (PAMPs) in response to bacterial virulence factors and epithelial stress, acting as a chemoattractant for neutrophils and contributing to sustained inflammation and tissue damage by promoting excessive neutrophil infiltration into the airway lumen^18,19^. IL-6, in contrast, is a pleiotropic cytokine involved in acute-phase responses, often upregulated following inflammatory signals, recognition of PAMPs, and other secondary pathways^18^. Elevated levels of IL-6 and IL-8 have been consistently associated with disease severity, frequent exacerbations, and accelerated lung function decline in patients with chronic airway conditions^20,21^.

The dual effects of mucin highlight a critical gap in our understanding of its influence on the outcome of bacterial infection and treatment. In particular, the effects of mucin-rich environments on modulating bacteriophage (phage), bacterial, and epithelial interactions remain largely undefined^22–24^. Phage therapy has re-emerged as a promising approach against multidrug-resistant bacteria due to its specificity, self-amplification, and minimal impact on commensal microbiota^25,26^. However, applying phages to mucus-dense tissues presents unique challenges, including limited diffusion, altered receptor accessibility, and potential immune interference^27–30^. The “Bacteriophage Adherence to Mucus” (BAM) model proposes that phages bind mucin glycans, increasing their concentration at mucosal surfaces and enhancing pathogen clearance^31–35^. Conversely, other studies suggest that mucins may act as a physical barrier, impeding phage diffusion and access to bacterial hosts^36–38^. In other bacterial models, such as *Flavobacterium columnare*, mucins have been shown to trigger bacterial protective stress responses or enhance adaptive mechanisms like CRISPR-Cas immunity^39^. These contrasting findings underscore the complexity of mucin-phage-bacteria interactions across different biological systems. Thus, the impact of mucins appears highly context-dependent, shaped by mucin type, microbial behavior, and infection dynamics^29,39,40^. In addition, little is known about how mucins influence bacterial evolution under phage pressure. Phage resistance commonly results in fitness costs or trade-offs in host bacteria, including altered motility, surface modifications, or reduced virulence^41–43^. Whether mucins mitigate or amplify these trade-offs, and how they shape the phenotypic and genotypic landscape of phage-resistant populations, remains to be determined.

In this study, we used a cell culture model to investigate how a mucin-enriched microenvironment modulates epithelial responses, *P. aeruginosa* virulence, phage efficacy, and evolution of phage resistance. We show that exogenous mucin induces CRD-like transcriptional reprogramming in epithelial cells, enhances bacterial cytotoxicity and inflammation, and selectively represses bacterial virulence pathways. Despite this, mucin synergizes with bacteriophage DMS3vir to protect epithelial cells in a bacterial load-dependent manner without eliciting antiviral immune responses. Moreover, mucin shapes the phenotypic and genetic profiles of phage-resistant *P. aeruginosa*, affecting virulent traits and resistance trade-offs. These findings provide new insights into the tripartite interactions between epithelium, bacteria, and phages, in mucin-rich conditions, with implications for optimizing phage therapies in mucosal environments.

## 2. Results

### 2.1. Mucin enrichment induces CRD-like transcriptional remodeling in airway epithelial cells

Mucus hypersecretion is a hallmark of chronic airway diseases such as COPD and CF^2,4^. To explore how a mucin-rich microenvironment influences epithelial gene expression, RNA sequencing was performed on A549 airway epithelial cells exposed to 0.2% porcine gastric mucin (PGM) for 18 h. Principal component analysis (PCA) revealed consistent intra-group clustering and clear separation between control and PGM-exposed groups, indicating a mucin-induced transcriptomic shift (Fig. 1a). Differential expression analysis identified 223 upregulated and 115 downregulated genes (log2 fold change > 0.5), and a stricter cutoff revealed 52 upregulated and 37 downregulated genes (log2 fold change > 1) in the PGM-exposed cells (Fig. 1b). The mucin-rich environment induced regulation of genes associated with stress and inflammation (response to hypoxia, unfolded protein, endoplasmic reticulum stress, tumor necrosis factor, oxidative stress), metabolic reprogramming (glycolytic process), cell adhesion, junction maintenance, epithelial to mesenchymal transition, and growth-factor signaling (EGF, TGF-β, and FGF) (Fig. 1c, d). These transcriptional changes are consistent with commonly observed in chronic lung disease^2,6,7^. Thus, exposure to exogenous mucin alone is sufficient to induce a CRD-like gene expression signature, offering a physiologically relevant platform to investigate epithelial responses and epithelial–pathogen interactions under mucosal conditions.

**Figure 1.**
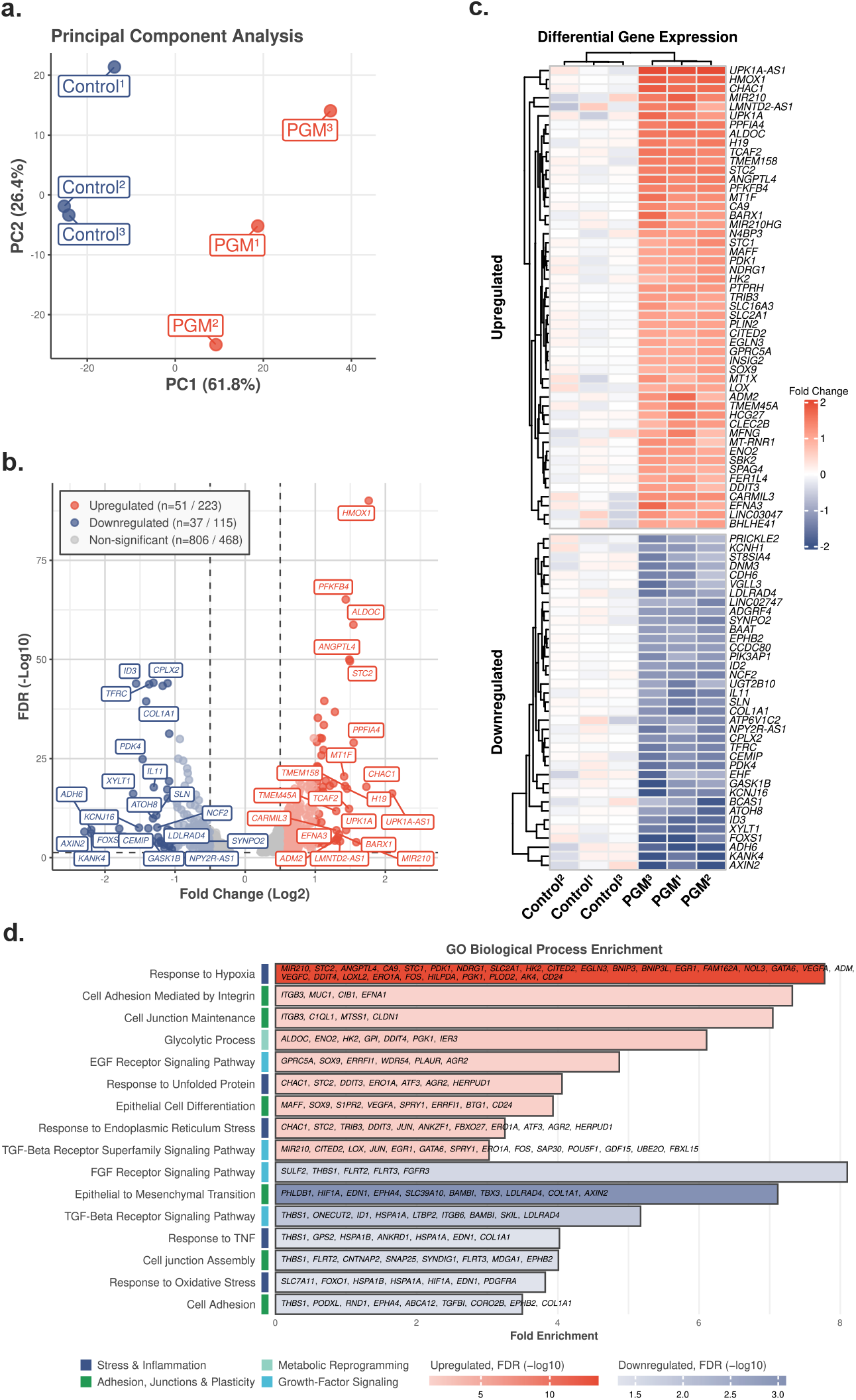
Transcriptomic analysis of A549 cells exposed to mucin rich environment. **(a)** Principal component analysis (PCA) of global gene expression in A549 cells incubated with 0.2% porcine gastric mucin (PGM) or control medium for 18 h (n = 3 per group). Each point represents a biological replicate. **(b)** Volcano plot of differential gene expression between PGM-treated and control cells. Log2 fold change (x-axis) is plotted against the –log10 FDR-adjusted p-value (y-axis). Significantly regulated genes were defined by FDR > 0.5, with log2 fold change > 1 or > 0.5 (upregulated, red), and log2 fold change > -1 or > -0.5 (downregulated, blue). **(c)** Heatmap of significantly differentially expressed genes between conditions (FDR < 0.05, log2 fold change > 1). Genes are hierarchically clustered and grouped into upregulated (top) and downregulated (bottom) sets. Columns represent biological replicates. Color scale indicates expression values. **(d)** Gene ontology of biological process enrichment analysis of upregulated (red) and downregulated (blue) genes. Bars represent significantly enriched biological processes, ranked by fold enrichment, with color intensity corresponding to statistical significance (–log10 FDR).

### 2.2. Mucin enriched environment enhances *Pseudomonas aeruginosa* cytotoxicity and amplifies epithelial inflammatory responses

To investigate how mucin-rich environment alters bacterial virulence, the cytotoxicity of *P. aeruginosa* was assessed with infection in A549 epithelial cells using bacterial strains PA14 and 573. PA14 exhibited significant cytotoxicity at 10^4^ CFUs, while strain 573 required a higher inoculum (10^6^ CFUs) to elicit similar effects (Fig. 2a). Pre-exposure of A549 cells with 0.2% PGM significantly enhanced cytotoxicity induced by both strains at 4 h post infection, indicating that mucin enrichment sensitizes epithelial cells to bacterial attack (Fig. 2b). This effect was independent of exposure timing with mucins, as both pre-incubation and co-incubation with PGM led to similar increases of epithelial cell death (Fig. 2c). A similar trend was observed in HT-29 intestinal epithelial cells, where PGM exposure increased susceptibility to PA14-induced cytotoxicity (Fig. 2d), suggesting that this phenomenon is not limited to airway epithelial cells. Notably, PGM alone had no detectable cytotoxic effect on either cell line, underscoring its role in modulating epithelial-bacteria interactions rather than decreasing cell survival. To explore the underlying mechanisms, bacterial internalization was quantified in mucin-exposed cells. Surprisingly, PGM significantly reduced *P. aeruginosa* uptake in both A549 and HT-29 cells (Fig. 2e, f), revealing an inverse relationship between cytotoxicity and bacterial internalization. These findings suggest that mucin does not enhance bacterial invasion, but rather exacerbates epithelial cells susceptibility, possibly by amplifying the impact of secreted virulence factors.

**Figure 2.**
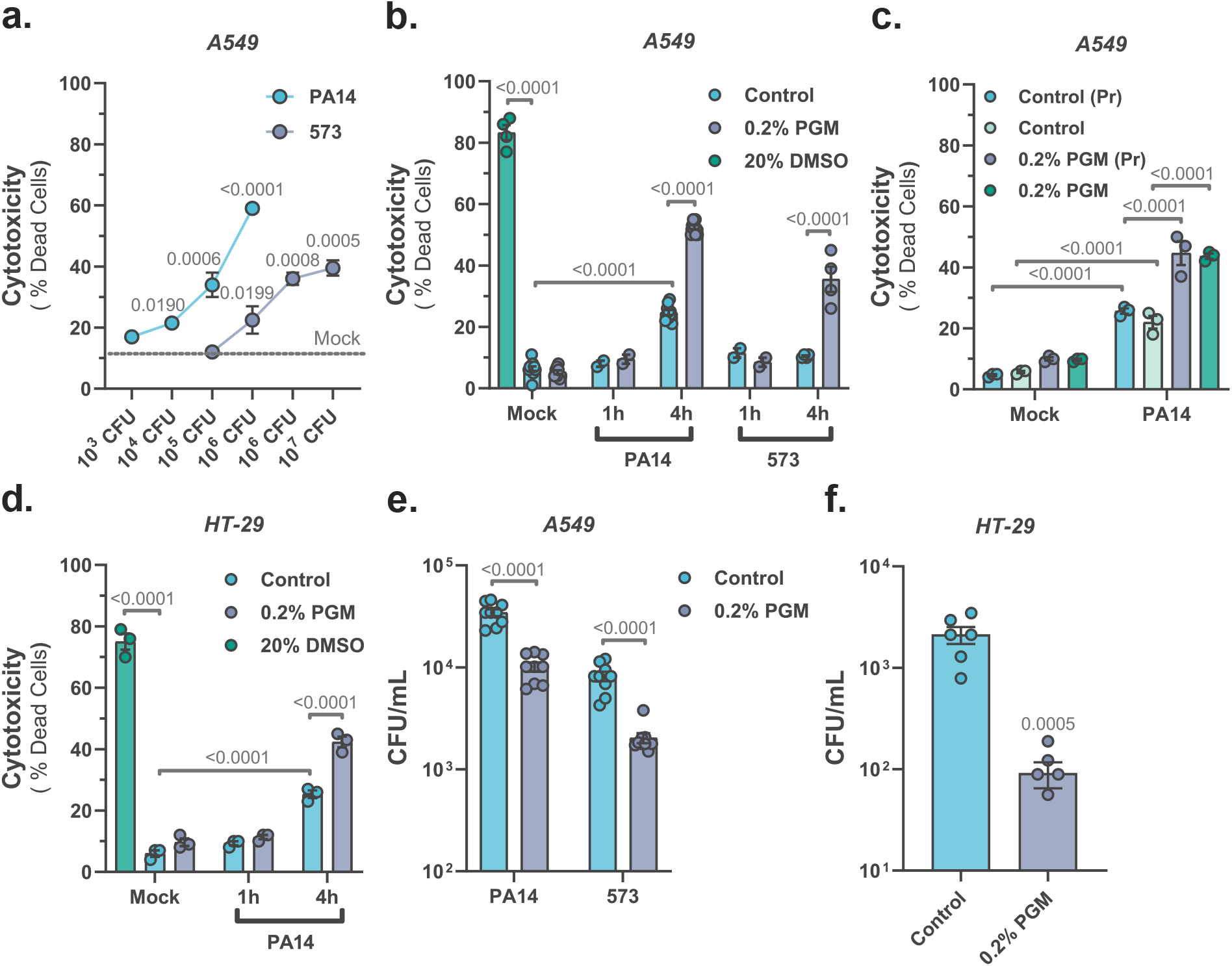
Mucin enrichment increases epithelial cytotoxicity and reduces bacterial internalization upon *P. aeruginosa* infection. **(a)** Bacterial cytotoxicity on A549 cells infected with *P. aeruginosa* PA14 and 573 strains for 4 h (n = 3). **(b)** Bacterial cytotoxicity on A549 cells pre-incubated with porcine gastric mucin (PGM) for 18 h prior to infection with PA14 or 573 (1 x 10^5^ CFU). **(c)** Bacterial cytotoxicity on A549 cells pre-exposed (Pr) to PGM or co-exposed to PGM and PA14 for 4 h. **(d)** Bacterial cytotoxicity on HT-29 cells pre-incubated with PGM prior PA14 infection (1 x 10^5^ CFU). 20% DMSO served as a positive control for cytotoxicity. **(e)** Bacterial internalization in A549 cells pre-incubated with PGM prior PA14 or 573 infections (1 x 10^5^ CFU) for 1 h. **(f)** Bacterial internalization in HT-29 cells pre-incubated with 0.2% PGM prior PA14 infection (1 x 10^5^ CFU) for 1 h. Quantitative data represent mean ± s.e.m. from at least two independent experiments. Statistical comparisons were performed using two-way ANOVA with multiple comparisons and Sidak’s post hoc test **(a-e)**, and unpaired two-tailed t-test **(f)**; p-values and replicate numbers are indicated.

Next, the impact of mucin-rich conditions on the expression of *IL6* and *IL8*, two key pro-inflammatory cytokines commonly elevated in CRD^9,10^, was assessed in A549 cells infected with *P. aeruginosa*. The *IL8* expression was induced after 4 h of PA14 infection and further amplified in the presence of PGM, particularly at later time point (Fig. 3a). Both PA14 and 573 strains triggered *IL8* and *IL6* expression under mucin-enriched conditions (Fig. 3b, c), while PGM alone did not induce cytokine expression, indicating that the inflammatory amplification is infection-dependent. These trends were also observed in HT-29 cells, where PGM exposure enhanced *IL8* expression during PA14 infection (Fig. 3d). The *IL6* expression remained undetectable in HT-29 cells across all tested conditions (data not shown), consistent with previous findings^44,45^. Collectively, these results suggest that mucin-rich environments sensitize epithelial cells to bacterial virulence, amplifying both cytotoxic and inflammatory response without promoting bacterial invasion.

**Figure 3.**
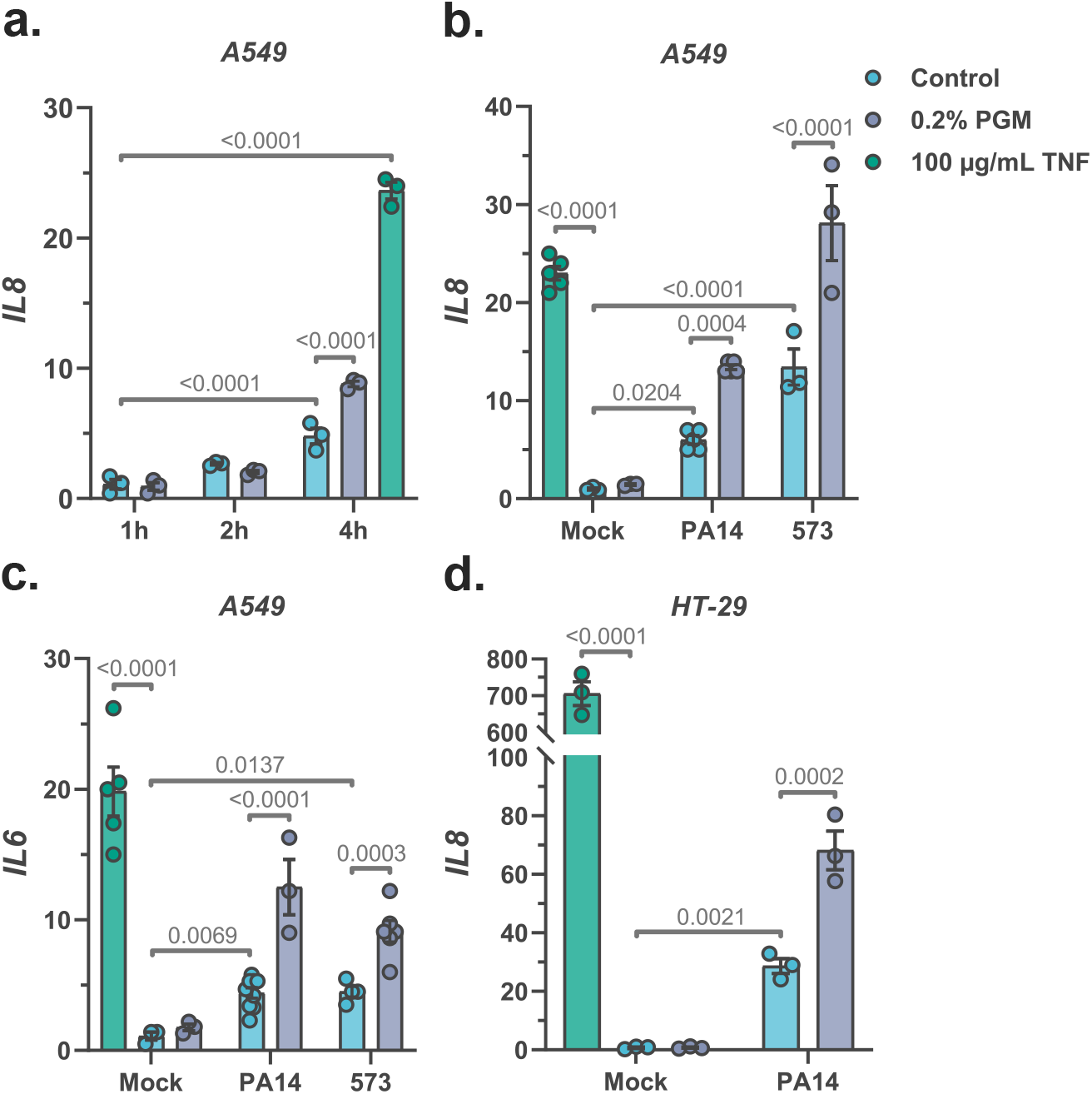
Mucin rich environment enhances pro-inflammatory gene expression during *P. aeruginosa* infection. **(a)** Time-course analysis of relative *IL8* mRNA expression in A549 cells pre-exposed to porcine gastric mucin (PGM) for 18 h prior *P. aeruginosa* PA14 infection (10^5^ CFU). **(b–c)** Relative *IL8* and *IL6* mRNA levels in A549 cells pre-incubated with PGM and infected with PA14 or 573 (1 x 10^5^ CFU) for 4 h. **(d)** *IL8* mRNA expression in HT-29 cells pre-incubated with PGM and infected with PA14 (1 x 10^5^ CFU) for 4 h. In all panels, cells were treated with 100 µg/mL TNF as a positive control. Quantitative data represent mean ± s.e.m. from at least two independent experiments. Statistical comparisons were performed using two-way ANOVA with multiple comparisons and Sidak’s post hoc test **(a-d)**; p-values and replicate numbers are indicated.

### 2.3. Mucin enrichment selectively decreases quorum sensing and virulence gene expression in *P. aeruginosa*

To evaluate whether mucin enrichment affects bacterial fitness, the growth of *P. aeruginosa* strains PA14 and 573 were monitored in the presence of 0.2% PGM. Growth curves showed no significant differences under PGM exposure (Fig. 4a, b). Additionally, neither strain was able to grow in nutrient-free condition supplemented PGM, confirming that PGM does not serve as a significant carbon or energy source (Fig. 4c, d). Although PGM did not influence bacterial growth, its impact on virulence gene regulation was notable. *P. aeruginosa* was exposed to PGM to determine whether mucin enriched condition alters bacterial virulence at the transcriptional level. Expression of quorum sensing (QS) and the virulence-associated gene *algD*, a key gene in alginate biosynthesis and biofilm formation, were quantified in both PA14 and 573 strains. Among QS genes, *rhlI* in PA14 and *rhlR* in 573 showed modest reductions, while expression of the other QS regulators remained unchanged (Fig. 4e, f). Notably, *lasR* and *lasI* transcripts were undetectable in strain 573 under all tested conditions, consistent with its reduced virulence phenotype (Fig. 4f). In both strains, exposure to PGM led to significant downregulation of *algD*, suggesting that mucin environment represses biofilm-associated pathways, rather than broadly suppressing QS. Pyocyanin production, a QS-regulated cytotoxin, was also evaluated. While overall levels varied over time, pyocyanin production by PA14 remained unaffected by PGM exposure (Fig. 4g). Taking together, these results indicate that while mucin-rich conditions do not impair bacterial proliferation, they can modulate transcriptional pathways associated with virulence. This underscores the complexity of epithelial-derived signals in modulating bacterial pathogenicity and highlights mucin as a context-dependent regulator of virulence gene expression.

**Figure 4.**
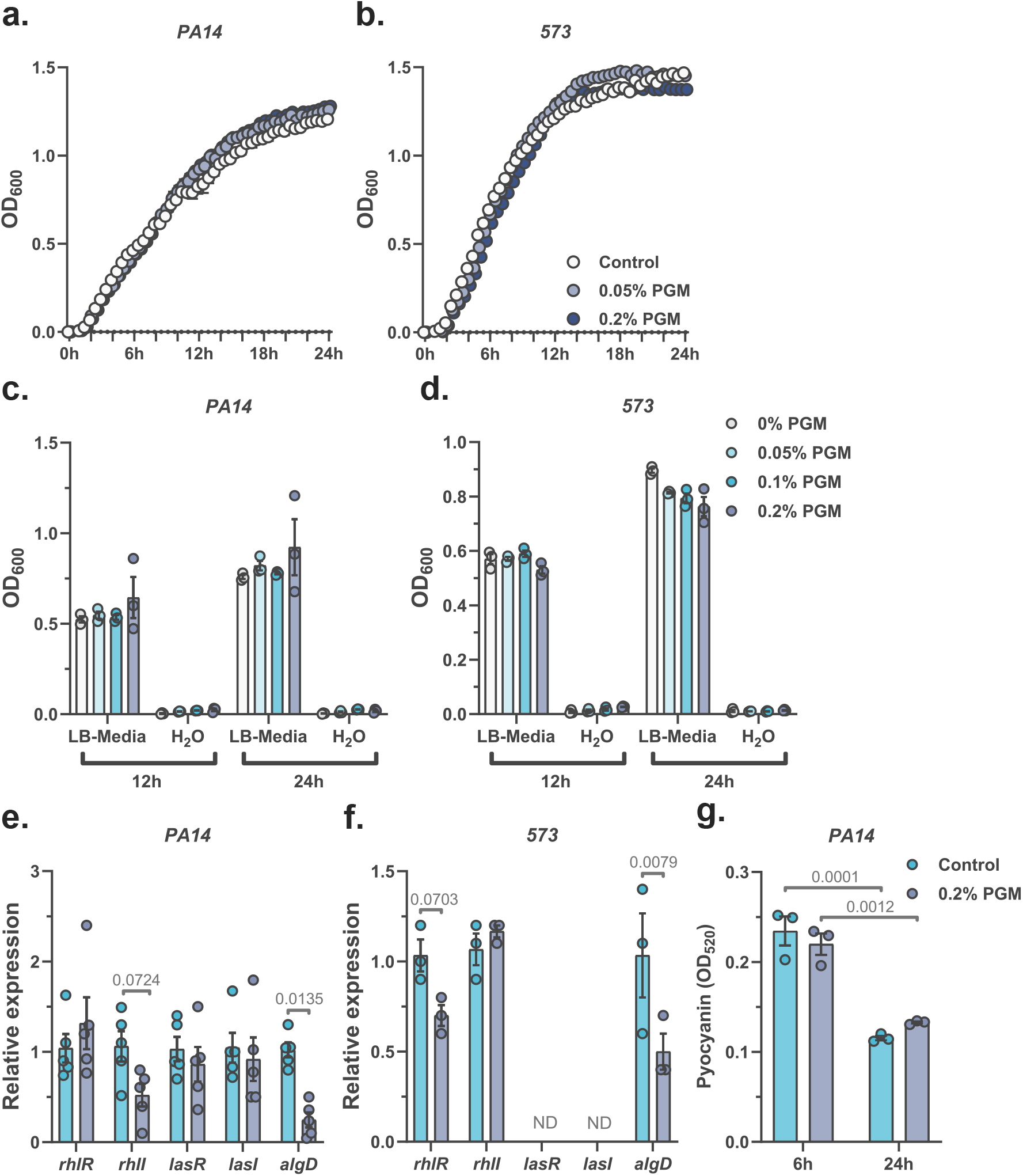
PGM does not serve as a nutrient source but modulates virulent gene expression in *P. aeruginosa*. **(a–b)** Bacterial growth curves of *P. aeruginosa* PA14 and 573 in LB medium supplemented with indicated porcine gastric mucin (PGM) concentration, measured by optical density at 600 nm (OD_600_) over 24 h (n = 5). **(c–d)** Endpoint OD_600_ readings of PA14 and 573 cultures incubated for 12 or 24 h in nutrient-free water (H_2_O) or LB media with increasing concentrations of PGM. **(e–f)** Relative expression of quorum sensing (QS) and virulence-associated genes in PA14 and 573 at 1 x 10^8^ CFU following 4 h incubation in LB supplemented with PGM. ND = not detected. **(g)** Pyocyanin production by PA14 (1 x 10^8^ CFU) in the presence of PGM. Values represent absorbance at 520 nm (OD_520_). Quantitative data represent mean ± s.e.m. from at least two independent experiments. Statistical comparisons were performed using two-way ANOVA with multiple comparisons and Sidak’s post hoc test **(a-g)**; p-values and replicate numbers are indicated.

### 2.4. Mucin-enriched environment enhances phage protection efficacy

To evaluate how mucin enrichment influences phage–bacterium interactions, the ability of DMS3vir phage to suppress *P. aeruginosa* PA14 growth were tested. DMS3vir effectively inhibited bacterial proliferation in a concentration-dependent manner, although regrowth was observed after approximately 18 h, consistent with emergence of phage-resistant clones (Fig. 5a). Notably, the presence of 0.2% PGM was unable to interfere with phage-mediated bacterial suppression (Fig. 5b).

**Figure 5.**
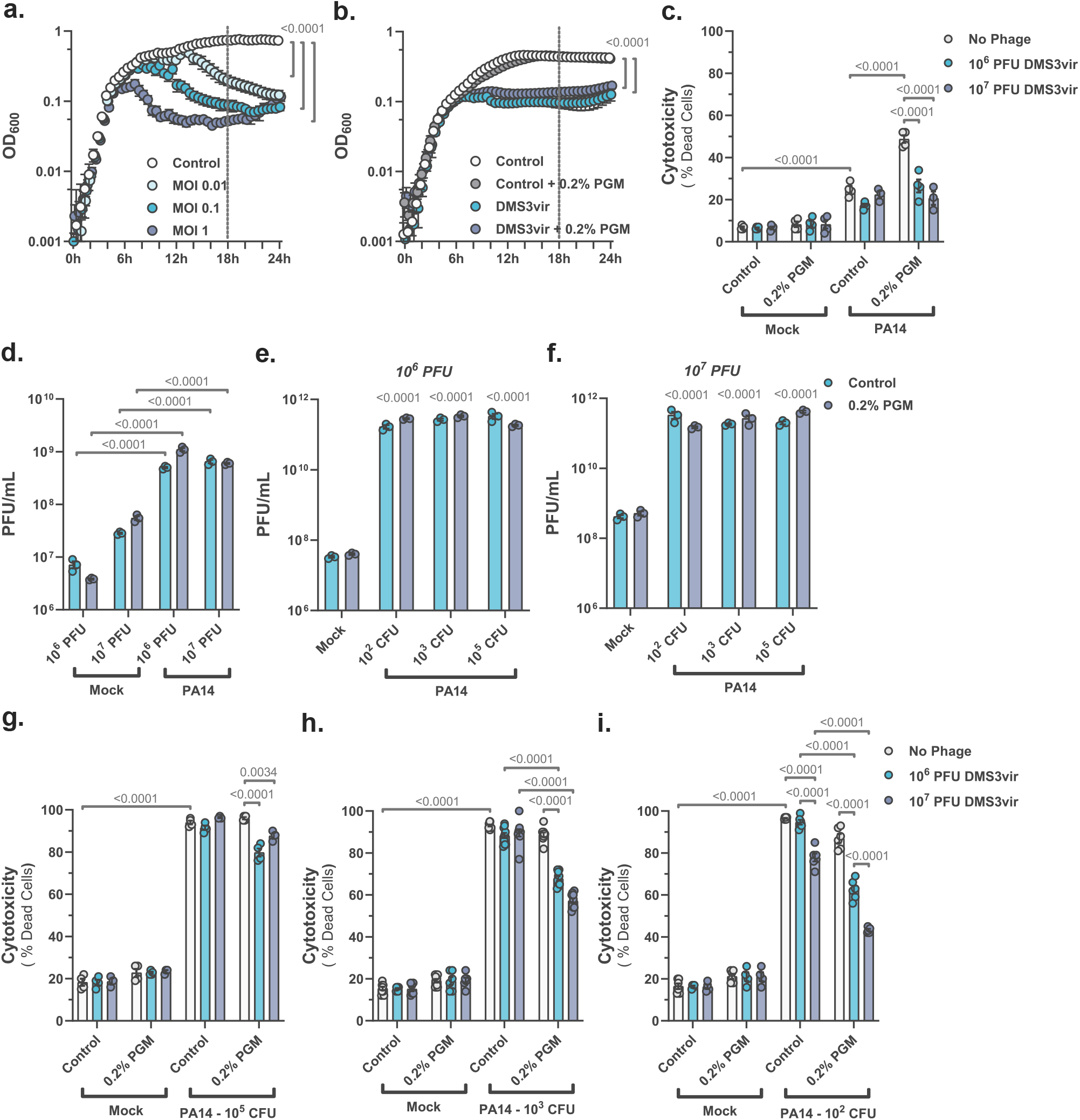
Mucin enrichment enhances the protective activity of phage DMS3vir against *P. aeruginosa* infection. **(a)** Bacterial growth curves of *P. aeruginosa* PA14 (1 x 10^5^ CFU) incubated with DMS3vir phage at different multiplicities of infection (MOIs) (n = 10). **(b)** Bacterial growth curves of PA14 (1 x 10^5^ CFU) co-exposed with DMS3vir phage (MOI 1) and porcine gastric mucin (PGM) (n = 5). **(c)** Bacterial cytotoxicity on A549 cells pre-incubated with DMS3vir and PGM for 18 h, followed by infection with PA14 (1 x 10^5^ CFU) for 4 h. **(d)** DMS3vir quantification of culture supernatants from the same experiment in **(c)**, collected after 4 h of infection. **(e–f)** Phage quantification from culture supernatants of A549 cells pre-exposed to PGM and DMS3vir prior PA14 infection at varying initial loads. **(g–i)** Bacterial cytotoxicity on A549 pre-exposed to PGM and DMS3vir, following PA14 infection at bacterial initial concentrations of **(g)** 1 x 10^5^, **(h)** 1 x 10^3^, or **(i)** 1 x 10^2^ CFU for 24 h. Quantitative data represent mean ± s.e.m. from at least two independent experiments. Statistical comparisons were performed using two-way ANOVA with multiple comparisons and Sidak’s post hoc test **(a-i)**; p-values and replicate numbers are indicated.

To assess whether mucin enrichment affects phage protection in epithelial cells, A549 were co-exposured to 0.2% PGM and DMS3vir prior to *P. aeruginosa* PA14 infection. After 4 h, mucin-enhanced cytotoxicity was significantly reversed by phage pre-incubation, restoring epithelial cell viability to levels similar to non-mucin conditions (Fig. 5c). Quantification of viral particles by titration revealed that DMS3vir phage replicated significantly in the presence of bacteria, and this replication was unaffected by PGM (Fig. 5d). Extended incubation of 24 h confirmed that phage amplification remained stable regardless of mucin exposure or initial bacterial load (Fig. 5e, f). Interestingly, 24 h co-culture revealed that mucin enhanced the protective effects of DMS3vir on the epithelial cells in a manner that depended on the initial bacterial inoculum. At high bacteria density (10^5^ CFUs), phage pre-incubation alone had limited impact, but when combined with PGM, epithelial cell viability improved modestly (Fig. 5g). In contrast, at lower bacterial inoculum (10^2^–10^3^ CFUs), co-incubation with phage and PGM synergistically preserved epithelial viability, indicating a threshold-dependent enhancement of phage protection by mucin (Fig. 5h, i).

Next, the influence of the phage on epithelial inflammatory responses in mucin-rich conditions was examined. As shown above, PGM alone amplified *IL6* and *IL8* expressions during PA14 infection (Fig. 6a, b). Interestingly, DMS3vir incubation selectively reduced *IL8* expression regardless of PGM presence, while *IL6* levels remained unaffected. Additionally, phage exposure did not trigger antiviral signaling pathways, as evidenced by unchanged *IFNB* expression (Fig. S1) and undetectable *IFNL1* transcripts (data not shown).

**Figure 6.**
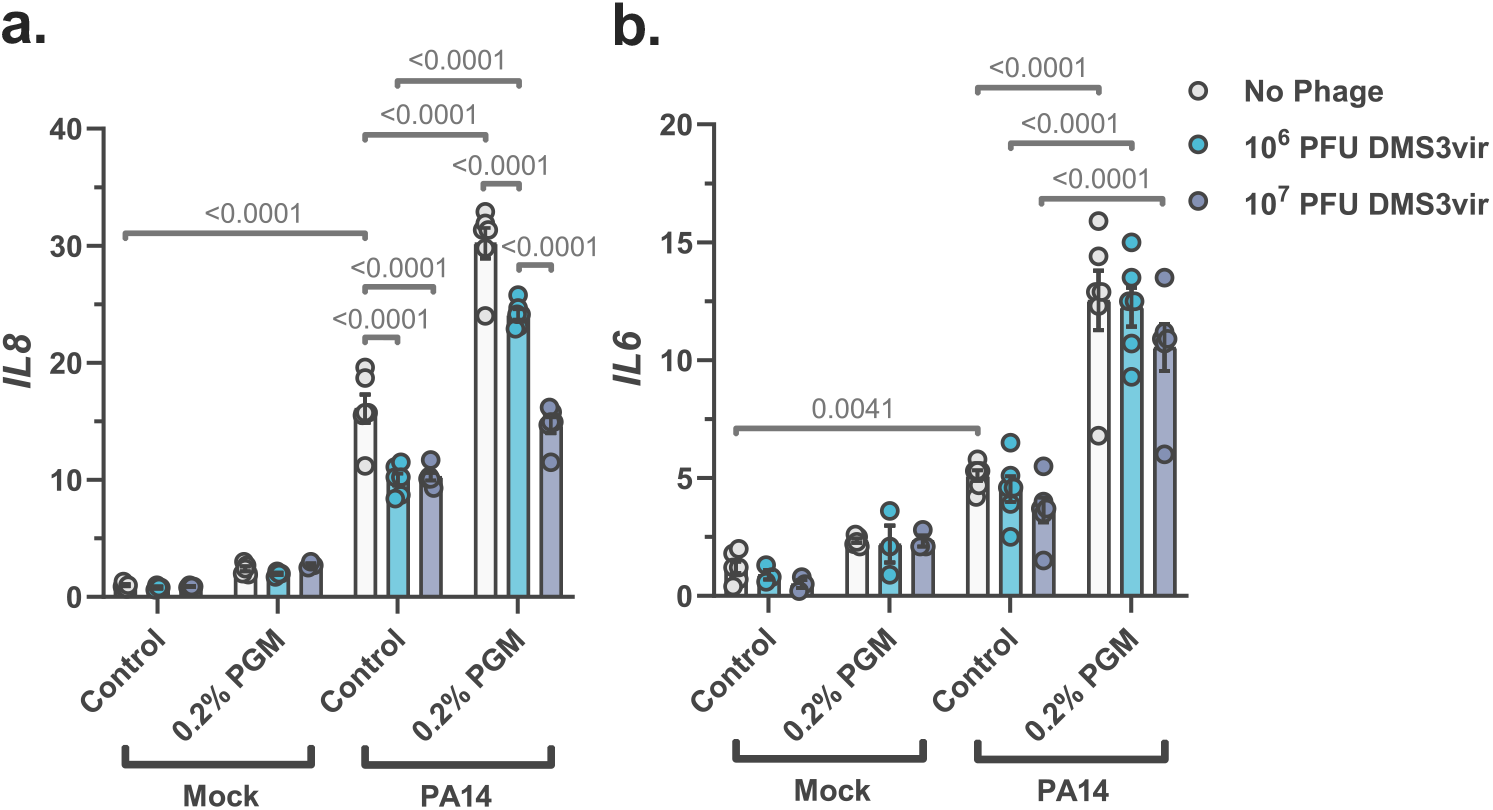
Phage DMS3vir selectively attenuates *IL8*, but not *IL6*, expression in *P. aeruginosa*-infected epithelial cells. **(a-b)** Relative expression of *IL8* and *IL6* mRNA of A549 cells pre-incubated with DMS3vir and porcine gastric mucin (PGM) for 18 h, followed by infection with *P. aeruginosa* PA14 (1 x 10^5^ CFU) for 4 h. Quantitative data represent mean ± s.e.m. from at least two independent experiments. Statistical comparisons were performed using two-way ANOVA with multiple comparisons and Sidak’s post hoc test **(a-b)**; p-values and replicate numbers are indicated.

These results suggest that mucin-rich microenvironments enhance the protective efficacy of DMS3vir phage while not impairing its replication, especially at submaximal bacterial loads relevant for real life infections. Moreover, DMS3vir exerts pathway-specific immunomodulatory effects without triggering antiviral responses, highlighting its compatibility with mucosal environments.

### 2.5. Mucin-enriched environment modulates phenotypic trade-offs in phage-resistant *P. aeruginosa*

To investigate how mucin exposure influences the evolution of phage-resistance, A549 cells were co-incubated with 0.2% PGM and DMS3vir phage prior *P. aeruginosa* PA14 infection. After 24 h, bacteriophage-insensitive mutants (BIMs) were isolated. Six independent clones were selected: three from standard conditions (BIM1-3) and three from mucin-enriched conditions (BIM(M)1-3). Control isolates not exposed to phage, including PA14(WT) (ancestral strain), PA14(C) (A549-exposed), and PA14(M) (A549 + PGM), retained their characteristic smooth, mucoid morphology with yellow-green pigmentation (Fig. 7a). By contrast, BIM clones developed smaller and rougher colonies with enhanced green pigmentation, suggestive of increased pyocyanin production (Fig. 7b). BIM(M) isolates displayed intermediate features, transitioning between rough and smooth morphologies with yellowish-green pigmentation, although BIM(M)1 isolate retained a rougher phenotype (Fig. 7c). Phage resistance was confirmed phenotypically, where DMS3vir effectively suppressed growth of control strains but failed to inhibit BIM or BIM(M) clones (Fig. 7d, e, f). Whole-genome sequencing revealed mutations across multiple functional categories, including virulence factors, surface modification, motility, surface attachment, metabolism, etc, (Fig. 7g, S2a). Notably, BIM(M) isolates exhibited a broader mutational spectrum, suggesting additional selective pressures in the mucin-rich environment. CRISPR spacer analysis confirmed that resistance was not mediated by spacer acquisition but likely by surface modification (Fig. S2c).

**Figure 7.**
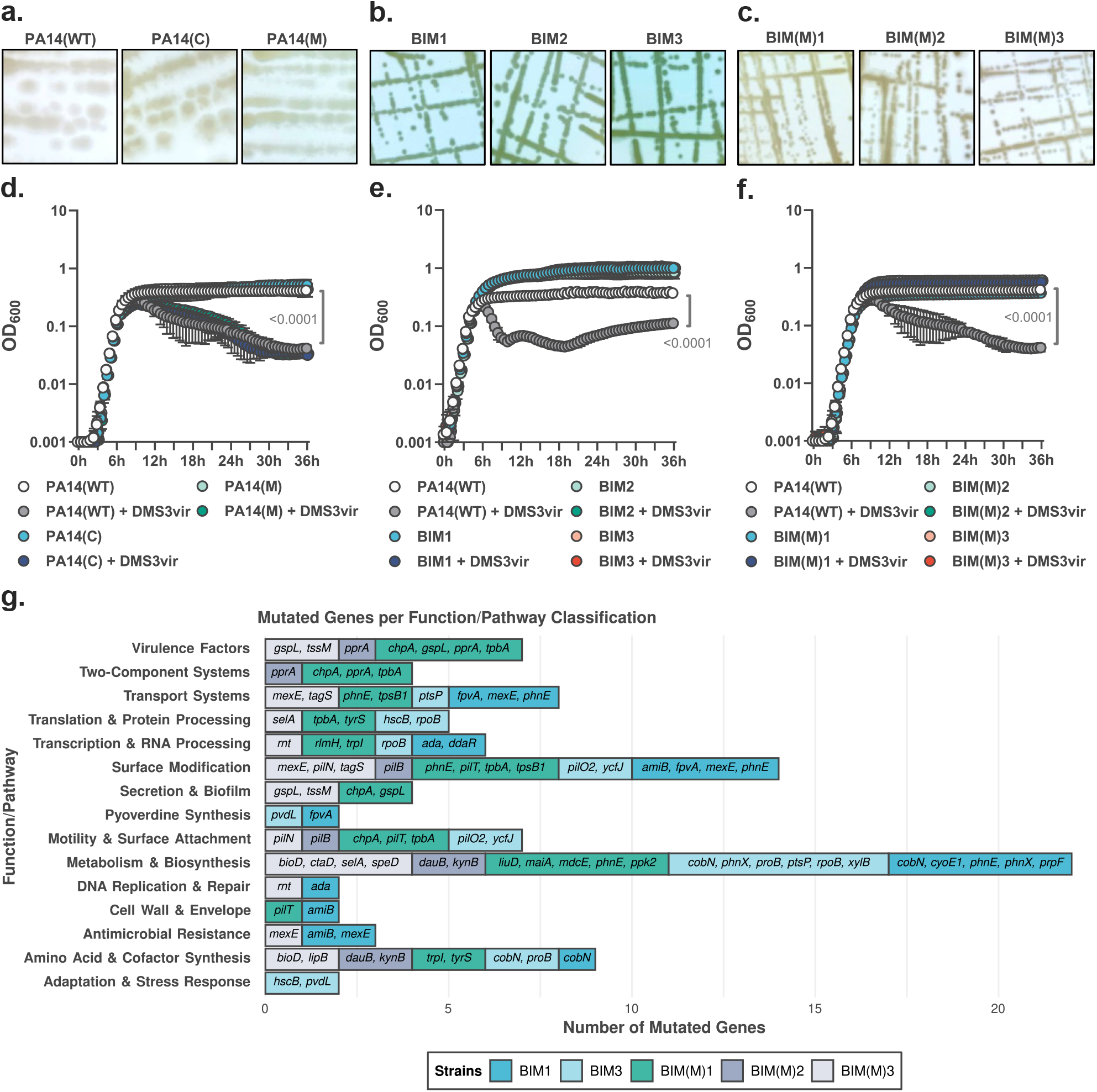
Isolation and genomic characterization of phage-resistant *P. aeruginosa* selected in the presence of epithelial cells and mucin rich environment. **(a)** Representative images of colony morphology of *P. aeruginosa* PA14 clones isolated after 24 h co-culture at 1 x 10^5^ CFU with A549 cells pre-exposed to 0.2% porcine gastric mucin (PGM) and DMS3vir phage (10^7^ PFU). **(a)** Control isolates: PA14(WT) (ancestral strain), PA14(C) (exposed to A549), PA14(M) (A549 + PGM). BIM isolates: **(b)** BIM1-2 (A549 + DMS3vir) and **(c)** BIM(M)1-3 (A549 + DMS3vir + PGM). Images show top-down views of individual (squared insert) and pool (circular insert) colonies. **(d-f)** Bacterial growth curves of **(d)** PA14 control clones, **(e)** BIM, and **(f)** BIM(M) exposed to DMS3vir (1 x 10^7^ PFU). Growth curves represent four replicates. **(g)** Functional/pathway annotation of whole genome sequenced genes mutated in BIM1-2 and BIM(M)1-3 strains. Mutated genes shared with PA14(WT), PA14(C), or PA14(M) were excluded. Bars indicate the number of mutated gene hits per category. Quantitative data represent mean ± s.e.m. from at least two independent experiments. Statistical comparisons were performed using two-way ANOVA with multiple comparisons and Sidak’s post hoc test **(d-f)**; p-values and replicate numbers are indicated.

Phenotypical assays revealed distinct trade-off associated with phage resistance. After 6 h, BIM(M) strains maintained pyocyanin production comparable to controls, whereas BIM strains showed significantly reduced levels (Fig. 8a). At 48 h, however, BIM strains increased pyocyanin production, while BIM(M) levels remained stable (Fig. 8b). Despite these differences, both BIM and BIM(M) strains exhibited impaired biofilm formation compared to controls (Fig. 8c), indicating that phage resistance compromises biofilm capacity regardless of mucin exposure. Motility assays further distinguished the two phage-resistant groups. Swarming motility (flagella-dependent) was absent in BIM strains, while BIM(M)2 and BIM(M)3 retained intermediate activity, and BIM(M)1 remained non-motile, similar to the BIM group (Fig. 8d). Twitching motility (pilus-dependent) was lost in all BIM and BIM(M) isolates (Fig. 8e). Moreover, cytotoxicity assays in A549 cells revealed significantly reduced virulence in phage-resistant clones. At 4 h post-infection, both BIM and BIM(M) strains induced minimal epithelial cell death, in sharp contrast to the cytotoxicity of control strains (Fig. 8f). This trend persisted at 8 h, where phage-resistant strains remained significantly less cytotoxic (Fig. 8g).

**Figure 8.**
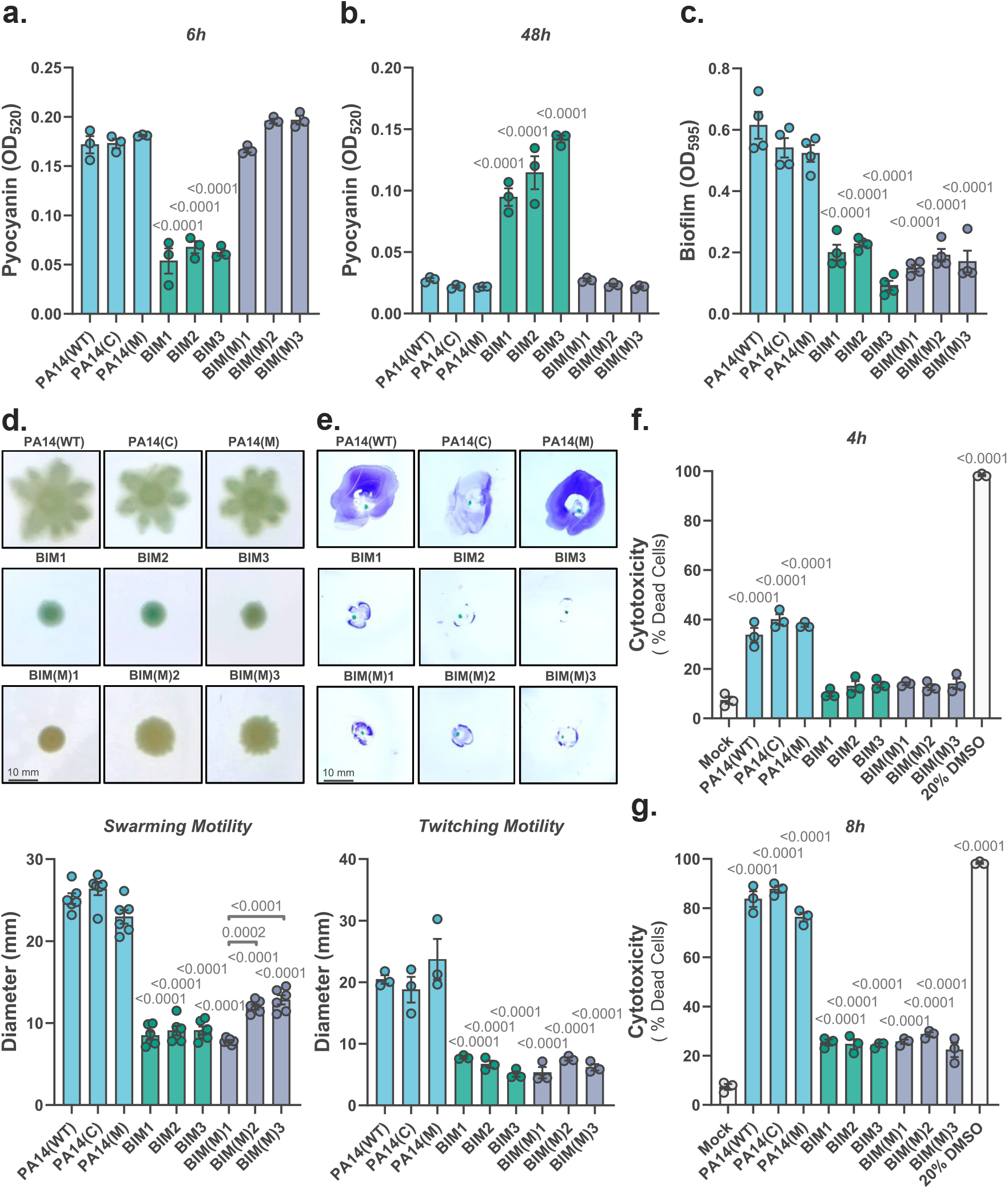
Phenotypic characterization of phage-resistant *P. aeruginosa* mutants reveals mucin-dependent modulation of virulence traits. **(a–b)** Pyocyanin production by PA14 clones (1 x 10^8^ CFU) after **(a)** 6 h or **(b)** 48 h incubation in LB medium. **(c)** Biofilm formation measured by crystal violet staining (OD_595_) after 24 h at 37 °C under static condition. **(d)** Swarming motility: representative images (upper panel) and motility diameters (lower panel) of PA14 clones spotted (2 µL, OD_600_ = 0.1) after 24 h at 37 °C. **(e)** Twitching motility: representative images (upper panel) and motility diameters (lower panel) after 48 h at 37°C, stained with crystal violet. **(f–g)** Cytotoxicity in A549 cells infected with PA14 clones (1 x 10^5^ CFU) for **(f)** 4 h or **(g)** 8 h. A 20% DMSO treatment served as a positive control for cytotoxicity. BIM and BIM(M) denote phage-resistant clones isolated in the absence or presence of porcine gastric mucin (PGM), respectively, as described in Figure 7. Quantitative data represent mean ± s.e.m. from at least two independent experiments. Statistical comparisons were performed using one-way ANOVA with Turkey’s post hoc test **(a-g)**; p-values and replicate numbers are indicated.

Together, these findings demonstrate that a mucin-enriched microenvironment does not prevent the emergence of phage resistance but reshapes the phenotypic and genotypic landscape of resistant populations. The BIM(M) strains retained pyocyanin levels and partially preserved the swarming motility compared to wild-type strain, while exhibiting reduced cytotoxicity, biofilm formation, and loss of twitching motility. These underscore the complex interplay between mucosal environments, bacterial adaptation, and phage-mediated selection, and highlight the importance of epithelial-derived factors in shaping the evolutionary outcomes relevant to phage therapy.

## 3. Discussion

Chronic respiratory diseases (CRD) such as COPD and CF are characterized by excessive mucin accumulation in the airways, profoundly reshaping host-pathogen dynamics^7,9,46^. Here we show that a mucin-enriched microenvironment reprograms epithelial responses, enhances *P. aeruginosa* cytotoxicity, and potentiates bacteriophage-mediated protection. Specifically, porcine gastric mucin (PGM) exposure induced CRD-like transcriptional remodeling in airway epithelial cells, sensitized them to bacterial damage, selectively modified *P. aeruginosa* virulence factor expression, and enhanced epithelial cell protection conferred by phage DMS3vir. These findings provide mechanistic insights into how mucin-rich conditions influence infection dynamics and therapy outcomes, with implications for mucosal-targeted interventions in chronic lung disease.

First, we aimed to understand how exogenous mucins influence epithelial cells. Transcriptomic profiling revealed that exogenous mucin drives hypoxia-associated metabolic reprogramming in A549 cells, with increased glycolytic enzymes, hypoxia-response factors, and stress-associated regulators. These patterns mirror CRD airway epithelium^3,5–7^, and suggest that mucin-rich environments simulate inflamed mucosal conditions. Effects on growth factor signaling and epithelial-mesenchymal transition point to epithelial stress and remodeling, potentially weakening barrier function and promoting infection susceptibility.

Next, we explored the interaction of bacterial and epithelial cells with and without exogenous mucins. Surprisingly, PGM increased *P. aeruginosa* cytotoxicity in airway and intestinal epithelial cells while reducing bacterial internalization. MUC1, a transmembrane mucin, is known to interact with *P. aeruginosa* flagellin, facilitating adhesion on epithelial cell^47,48^. It is possible that the mucin-rich environments block MUC1 receptors, decreasing bacterial adhesion but increasing epithelial vulnerability to secreted virulence factors. Elevated *IL6* and *IL8* expression under mucin exposure supports heightened epithelial sensitivity, consistent with inflammatory exacerbation during mucosal infections^17,20,49^. Notably, IL-8 upregulation may amplify neutrophil-driven tissue damage, fueling a detrimental cycle of inflammation and remodeling^50^. Moreover, recent study linking flagellar motility to aggregation-mediated antibiotic tolerance in mucin environments^16^ further underscore the role of mucins in modulating bacterial access and epithelial host responses.

When looking at the influence of mucins on bacterial cells alone, we observed that PGM selectively downregulated virulence-associated genes, including *algD* and specific quorum sensing regulators, without impairing growth or pyocyanin production. This suggests mucin functions as a signaling modulator, tuning bacterial programs rather than serving as a nutrient source^14,40,51^, consistent with earlier findings that neither PGM nor purified MUC5B support *P. aeruginosa* growth^32,51^. While PGM contains MUC5AC with human-like glycosylation, its composition differs from human airway mucins, which also include MUC5B and non-mucin proteins^39,52–54^. These compositional differences may engage distinct bacterial sensing pathways and explain differential regulatory effects. Reduced *algD* expression, despite sustained growth, could weaken biofilm integrity and persistence^55,56^.

Since mucosal surfaces host a diverse community of bacteriophages, and phages could be used in treatment and prevention of mucosal infections, we looked into the tripartite interactions between phages, their bacterial host and epithelial cells. Strikingly, mucin-rich environment enhanced phage infection efficiency. Although PGM did not affect DMS3vir replication or lytic activity, the co-incubation significantly improved epithelial protection, particularly at lower bacterial loads. This threshold-dependent synergy suggests mucin alters the epithelial environment to favor phage-mediated defense. Such effects may define therapeutic windows during early infection or low-density colonization^35,38^. The possibility of combining mucin and phage for preventive strategies, such as mucosal colonization in high-risk individuals^30,57^, is reinforced by recent inhaled phage trials in CF patients that demonstrated reduced *P. aeruginosa* burden without adverse events^57^.

Phage treatment also modulated epithelial immune responses. DMS3vir reduced *IL8* expression during infection without affecting *IL6* or triggering type I/III interferon responses. Both cytokines are regulated by NF-κB and MAPKs, but *IL6* transcription is further controlled by cAMP and prostaglandin-responsive elements absent in *IL8*^19,58,59^. The selective IL-8 suppression may dampen neutrophil-driven inflammation while preserving other epithelial defenses, contributing to tissue protection in mucosal infections.

Understanding the influence of environment on evolution of phage resistance is central for phage-bacterium interactions and phage therapy. Here, phage-resistant *P. aeruginosa* clones displayed distinct adaptation patterns under mucin influence. While resistance uniformly emerged without CRISPR spacer acquisition, implicating surface modification as the dominant mechanism^60–62^, mucin-exposed BIM(M) isolates retained pigment production and partial motility compared with BIMs. Whole-genome sequencing revealed mutations in surface structure, motility, and secretion pathways, with broader diversity in BIM(M) strains, underscoring mucin’s role in shaping resistance trajectories^31,32,39,55,56^.

Swarming motility, driven by flagella, was completely lost in BIMs and in one BIM(M) isolate, whereas two BIM(M) clones retained partial swarming, suggesting that mucin preserves some motility functions under selective pressure. In contrast, twitching motility, dependent on type IV pili, was uniformly abolished across resistant clones, highlighting a broader fitness cost. Such motility impairments may restrict bacterial dissemination in mucosal niches and thereby enhance phage efficacy^15^. Despite differences in colony morphology and pigment production, all resistant clones showed reduced biofilm formation and attenuated cytotoxicity, consistent with a trade-off between resistance and virulence. However, BIM(M) isolates maintained pyocyanin and partial motility, suggesting that mucin tempers resistance costs and allows retention of virulence traits. Delayed pyocyanin accumulation in BIMs may reflect altered quorum sensing or metabolic adaptation^63–65^. Although based on limited isolates, these results highlight the environmental context in shaping bacterial evolution under phage pressure and permit larger-scale studies.

While our *in vitro* system recapitulates key CRD-like mucosal features, it lacks the cellular and immune complexity of the airway^66^. Future studies incorporating organoids, airway-on-chip systems, or animal models, alongside diverse phage cocktails, will be critical to fully capture mucin-bacterial-phage-epithelium interactions. Strategies that enhance phage delivery in mucinous environments or directly modulate mucin composition may represent promising therapeutic avenues^57^, including mucoactive-phage combinations^38^.

In summary, our findings reveal the dual role of mucin-rich environments: sensitizing epithelial cells to bacterial virulence while enhancing phage-mediated protection. By reshaping epithelial and microbial responses, mucin modulates both infection severity and therapeutic potential. Understanding these interactions lays the foundation for phage-based interventions tailored to mucosal surfaces in chronic airway disease.

## 4. Methods

### 4.1. Cell culture and reagents

A549 (ATCC®, CCL-185™) and HT-29 (ATCC®, HTB-38™) cells were maintained at 37 °C with 5% CO_2_ in DMEM (ThermoFisher, 25100039) supplemented with 10% fetal bovine serum (FBS; ThermoFisher, A5256701), 1% Glutamax (ThermoFisher, 35050038), and 1 U/mL penicillin/streptomycin (ThermoFisher, 15140122). Both cell lines were tested monthly for mycoplasma contamination by real-time PCR (primers Myc_526_F: ACATTGGGACTGAGATACGGC; Myc_1031_R: TCGTTTACGGCGTGGACTAC) using the *p_m16S* plasmid (addgene, 160549) as a positive control. For experiments, 1 x 10^5^ A549 or 2 x 10^5^ HT-29 cells (passages 10-25) were seeded in 96-well plates.

The mucin-enriched conditions were generated by supplementing culture medium with porcine gastric mucin (PGM; Sigma-Aldrich, M1778), a mucin source replicating structural features of human MUC5AC and previously validated in phage-bacteria-mucin studies^30,32,39,40,51,53,56^. A 2% (wt/vol) stock solution was prepared in PBS, autoclaved, filtered through 5 µm filters (Sigma-Aldrich, SLSV025LS), and diluted into medium prior use. PBS-only controls maintained equivalent volumes. Human TNF (100 ng/mL; ThermoFisher, PHC3011) served as a positive control for qPCR, and 20% DMSO (Avantor, 67-68-5) as a positive control for cell viability assays.

### 4.2. Bacterial strains and growth conditions

*P. aeruginosa* strain PA14 (DSM 19882), a highly virulent isolate, and strain 573, a less virulent clinical isolate from bone marrow interstitial fluid^32,67,68^, were cultured in Luria-Bertani Lennox medium (LB). For experiments, single colonies were grown overnight in 5 mL LB at 37 °C with agitation (200 rpm). Cultures were diluted 1:1 and incubated for 1 h under the same conditions, harvested by centrifugation (4,000 × *g*, 5 min, RT ∼22 °C), washed in PBS, and resuspended in LB or DMEM (10% FBS, 1% Glutamax, antibiotic-free). Suspensions were standardized to OD_600_ 0.3 (∼2 x 10^8^ CFU/mL) before dilution. Bacterial quantification was performed by CFU plating or OD_600_ readings (Multiskan FC, ThermoScientific). Growth was monitored at 600 nm (Tecan Intinite 220 PRO M Nano) every 30 min at 37 °C with shaking.

### 4.3. Cell viability

To assess bacterial cytotoxicity, culture supernatants and cells were collected into microtubes. Residual adherent cells were detached with 0.05% trypsin (ThermoFisher, 15400054; 5 min, 37 °C) and pooled. Samples were centrifuged (270 x *g*, 5 min, 4 °C), resuspended in DMEM (10% FBS), mixed 1:1 with Trypan Blue (ThermoFisher, 15250061), and transferred to cell counting chambers (Bio-Rad, 1450011). Viability was determined using TC20 automated cell counter (Bio-Rad).

### 4.4. Real-Time PCR

Total RNA was extracted using RNeasy kit (Qiagen, 74106). Reverse transcription performed with the High-Capacity cDNA Kit (ThermoFisher, 4368813) using 200 ng RNA. qPCR was conducted with Gotaq qPCR Master Mix (Promega, A6002) on a CFX96 C1000 cycler (Bio-Rad). Reactions contained 2 µL 1:10 diluted cDNA, 0.2 µM primers, and 10 µL total volume. Human 18S served as the reference gene, and *oprL* was used for *P. aeruginosa*. Data were analyzed in CFX Maestro v2.3 (Bio-Rad) using the 2^-ΔΔCT^ method.

### 4.5. Bacterial internalization

Epithelial cells were incubated with 400 µg/mL gentamicin (ThermoFisher, 15710064) for 1 h at 37 °C to eliminate extracellular bacteria (see Fig. S3). Following PBS washes, intracellular bacteria were released by treatment with 0.05% trypsin plus 0.2% Triton X-100 (Sigma-Aldrich, X100). CFU quantification was performed by serial dilutions and plating. Control groups without epithelial cells served as negative controls.

### 4.6. Pyocyanin production

Pyocyanin production was quantified using chloroform-HCl extraction. Overnight cultures were diluted to 2 x 10^8^ CFU/well in LB and incubated at 37 °C for 6 and 48 h in a 12-well plate (DTS-4 shaker, ELMI). Cultures were centrifuged (5000 x *g*, 10 min), and supernatants were mixed 5:3 with chloroform by repeated vortexing. The organic layer was mixed 2:1 with 0.2 M HCl, vortexed, and the aqueous phase absorbance was measured at 520 nm (Tecan).

### 4.7. Biofilm formation

Overnight cultures were diluted to OD_600_ = 0.1 and seeded in 96-well plates. Plates were incubated at 37 °C for 24 h under static conditions, washed with water, and stained with 0.1% crystal violet (20 min, RT). Excess stains were removed by washing, plates were dried, and biofilms were solubilized with 96% ethanol. Absorbance was read at 595 nm (Multiskan FC).

### 4.8. Bacteriophage

The lytic phage DMS3vir^69^ was propagated on *P. aeruginosa* strain PA14. Lysates were filtered (0.22 µm; Sarstedt, 83.1826.001), sterilized with 10% chloroform (Sigma-Aldrich, C2432), concentrated using 100 KDa Amicon filters (Sigma-Aldrich, UFC100), and purified with Endotrap HD 1/1 Kit (Lionex, LET0009). Endotoxin levels decreased from 7158 EU/10^9^ PFU to 2 EU/10^9^ PFU (Pierce Chromogenic Endotoxin Quant Kit, Thermo Fisher, A39552). Phage titers were determined by double-layer agar assay and stored at 4 °C.

### 4.9. Isolation of phage-resistant bacteria

Resistant clones were isolated from A549 cultures incubated with DMS3vir for 24 h. Triplicate colonies were obtained from two conditions: (i) BIM (A549 + DMS3vir) and (ii) BIM(M) (A549 + DMS3vir + 0.2% PGM). Control clones included (i) PA14(WT), (ii) PA14(C) (A549-exposed), and (iii) PA14(M) (A549 + PGM). Colonies were re-streaked ≥ 3 times for purification and stored at –80 °C in 15% glycerol.

### 4.10. Swarming motility

Cultures were diluted to OD_600_ = 0.1, and 2 µL was spotted onto LB soft agar. Plates were incubated at 37 °C for 24 h under static conditions. Swarming patterns were imaged alongside a ruler, and colony diameters measured using ImageJ2 v2.

### 4.11. Twitching motility

Bacterial colonies were stabbed to the bottom of LB agar plates with inoculation needle and incubated at 37 °C for 48 h. Agar was removed, plates were stained with 0.1% crystal violet, and twitching diameters were imaged and quantified with ImageJ2 v2.

### 4.12. RNA sequencing and analysis

Total RNA was extracted from A549 cells (n=3 per group; control and PGM-treated) with RNeasy Mini Kit. RNA integrity was confirmed (Agilent 2100 Bioanalyzer RIN>8). Strand-specific mRNA libraries were commercially sequenced on the DNBseq platform (BGI Tech, PE150, ∼20–30M reads/sample) in BGI, China. Processing followed the Chipster RNA-seq workflow (accessed October 2025; https://chipster.csc.fi/manual/rna-seq-summary.html) with defaut parameters. Differentially expressed genes considered significant (FDR < 0.05) in both DESeq2 (v.48.2) and edgeR (v4.6.3) tools were retained as the intersect (“Venn” intersection). Gene annotations used biomaRt (v2.62.1) and merged with org.Hs.eg.db (v3.30.0). PCA was performed on the top 1,000 variable genes. Visualization included ggplot2 (tidyverse v3.0.0) and ComplexHeatmap (v2.22.0). GO enrichment (clusterProfiler v4.6.2, org.Hs.eg.db) was curated and displayed as bar plots. Additional description available in Supplementary Information.

### 4.13. Whole-genome sequencing and analysis

Genomic DNA was extracted with DNeasy Blood & Tissue Kit (Qiagen, 69504) and quantified on Qubit fluorometer (Invitrogen). Libraries (≤800 bp inserts) were sequenced on DNBseq (BGI Tech, PE150). BIM2 sample sequencing failed due to technical issues. Reads were filtered with fastp (v0.24.0) and summarized with MultiQC (v1.31). Alignments used BWA-MEM (v0.7.19) and were processed with SAMtools (v1.21) and GATK MarkDuplicates (v4.6.2). Variants were called with SAMtools mpileup and VarScan (v2.4.2), annotated in R (VariantAnnotation v1.52.0, PA14 GFF and UniProt), and filtered to exclude synonymous and background variants. Heatmaps were generated with ComplexHeatmap. Genes were grouped by KEGG pathways/function and visualized as stacked bar plots. Assemblies were generated with SPAdes (v4.0.0), and CRISPR arrays were detected with minced (v0.2.0), parsed in R, and visualized as tile plots. Additional description available in Supplementary Information.

### 4.14. Statistical analysis

Statistical analyses were performed in GraphPad Prism v10.3.0. Two-tailed Student’s *t*-tests, one-way ANOVA with Turkey’s post hoc test, and two-way ANOVA with Sidak’s correction were applied as appropriate. Differential expression analyses used Wald tests (DESeq2) and quasi-likelihood F-tests (edgeR), with Benjamini-Hochberg correction. Exact *P*-values are reported. RNA-seq sample sizes were based on detecting fold-changes >1.5 with 80% power at α = 0.05. No statistical tests were applied to CRISPR spacer acquisition, results are shown descriptively. Variant calling results were not corrected for multiple testing due to low variant counts.

## 5. Data availability

The datasets generated during the current study are available in Supplementary Information files and at Zenodo (DOI: 10.5281/zenodo.17278999). The RNA sequencing data has been deposited in the NCBI Gene Expression Omnibus (GEO) under accession number GSE307737. Raw RNA from epithelial cells and bacterial whole-genome sequence reads are available in the NCBI Sequence Read Archive (SRA) under BioProject PRJNA1308451.

## 6. Code availability

All analysis scripts and visualization code are deposited in Zenodo (DOI: 10.5281/zenodo.17278999).

## 8. Acknowledgments

We would like to thank MSc Jenne Auramo and Mr Petri Papponen for help in the laboratory, Dr. Nina Chanisvili (George Eliava Institute of Bacteriophages, Microbiology & Virology, Tbilisi, Georgia) for donating *P. aeruginosa* strain 573, and Prof. Varpu Marjomäki for providing equipment and for donating reagents and the enterovirus *Coxsackievirus A9* (CVA9) used as a *IFNB* expression positive control. This study was funded by the Research Council of Finland grant (#346772) (L.-R.S.) and Centre for New Antibacterial Strategies (CANS) of the Arctic University of Norway (project ID #2520855) (G.M.F.A).

## 9. Author information

### Contributions

All authors contributed to the preparation of the manuscript. L.R.S., G.M.F.A, and D.O.P. contributed with project conception and design. L.R.S. and D.O.P. designed methodology. D.O.P., K.N., V.S, and L.W.G contributed with data acquisition and analysis. D.O.P. performed statistical analysis and bioinformatics. L.R.S. acquired and administered funding. All authors gave final approval for publication.

## References

1. Hauber, H.-P., Foley, S. C. & Hamid, Q. Mucin overproduction in chronic inflammatory lung disease. Can Respir J 13, 327–35 (2006).

2. Shah, B. K., Singh, B., Wang, Y., Xie, S. & Wang, C. Mucus Hypersecretion in Chronic Obstructive Pulmonary Disease and Its Treatment. Mediators Inflamm 2023, 8840594 (2023).

3. Symmes, B. A., Stefanski, A. L., Magin, C. M. & Evans, C. M. Role of mucins in lung homeostasis: regulated expression and biosynthesis in health and disease. Biochem Soc Trans 46, 707–719 (2018).

4. Fenker, D. E. et al. A Comparison between Two Pathophysiologically Different yet Microbiologically Similar Lung Diseases: Cystic Fibrosis and Chronic Obstructive Pulmonary Disease. Int J Respir Pulm Med 5, (2018).

5. Mikami, Y. et al. Chronic airway epithelial hypoxia exacerbates injury in muco-obstructive lung disease through mucus hyperconcentration. Sci Transl Med 15, (2023).

6. Worlitzsch, D. et al. Effects of reduced mucus oxygen concentration in airway Pseudomonas infections of cystic fibrosis patients. J Clin Invest 109, 317–25 (2002).

7. Montgomery, S. T., Mall, M. A., Kicic, A. & Stick, S. M. Hypoxia and sterile inflammation in cystic fibrosis airways: mechanisms and potential therapies. European Respiratory Journal 49, 1600903 (2017).

8. Pincikova, T. et al. Expression Levels of MUC5AC and MUC5B in Airway Goblet Cells Are Associated with Traits of COPD and Progression of Chronic Airflow Limitation. Int J Mol Sci 25, (2024).

9. Bals, R., Weiner, D. J. & Wilson, J. M. The innate immune system in cystic fibrosis lung disease. J Clin Invest 103, 303–7 (1999).

10. Hiemstra, P. S., McCray, P. B. & Bals, R. The innate immune function of airway epithelial cells in inflammatory lung disease. Eur Respir J 45, 1150–62 (2015).

11. Malhotra, S., Limoli, D. H., English, A. E., Parsek, M. R. & Wozniak, D. J. Mixed Communities of Mucoid and Nonmucoid Pseudomonas aeruginosa Exhibit Enhanced Resistance to Host Antimicrobials. mBio 9, (2018).

12. Moradali, M. F., Ghods, S. & Rehm, B. H. A. Pseudomonas aeruginosa Lifestyle: A Paradigm for Adaptation, Survival, and Persistence. Front Cell Infect Microbiol 7, 39 (2017).

13. Brockhausen, I., Falconer, D. & Sara, S. Relationships between bacteria and the mucus layer. Carbohydr Res 546, 109309 (2024).

14. Wheeler, K. M. et al. Mucin glycans attenuate the virulence of Pseudomonas aeruginosa in infection. Nat Microbiol 4, 2146–2154 (2019).

15. Greenwald, M. A. et al. Mucus polymer concentration and in vivo adaptation converge to define the antibiotic response of Pseudomonas aeruginosa during chronic lung infection. mBio 15, e0345123 (2024).

16. Higgs, M. G. et al. Flagellar motility and the mucus environment influence aggregation-mediated antibiotic tolerance of Pseudomonas aeruginosa in chronic lung infection. mBio 16, e0083125 (2025).

17. Barnes, P. J., Shapiro, S. D. & Pauwels, R. A. Chronic obstructive pulmonary disease: molecular and cellularmechanisms. European Respiratory Journal 22, 672–688 (2003).

18. Grebenciucova, E. & VanHaerents, S. Interleukin 6: at the interface of human health and disease. Front Immunol 14, (2023).

19. Cronin, J. G., Kanamarlapudi, V., Thornton, C. A. & Sheldon, I. M. Signal transducer and activator of transcription-3 licenses Toll-like receptor 4-dependent interleukin (IL)-6 and IL-8 production via IL-6 receptor-positive feedback in endometrial cells. Mucosal Immunol 9, 1125–1136 (2016).

20. Eidelman, O. et al. Control of the Proinflammatory State in Cystic Fibrosis Lung Epithelial Cells by Genes from the TNF-αR/NFκB Pathway. Molecular Medicine 7, 523–534 (2001).

21. Nakamoto, K. et al. Pseudomonas aeruginosa-derived flagellin stimulates IL-6 and IL-8 production in human bronchial epithelial cells: A potential mechanism for progression and exacerbation of COPD. Exp Lung Res 45, 255–266 (2019).

22. Carroll-Portillo, A. & Lin, H. C. Exploring Mucin as Adjunct to Phage Therapy. Microorganisms 9, (2021).

23. Pirnay, J.-P. et al. Personalized bacteriophage therapy outcomes for 100 consecutive cases: a multicentre, multinational, retrospective observational study. Nat Microbiol 9, 1434–1453 (2024).

24. Rodriguez-Gonzalez, R. A., Balacheff, Q., Debarbieux, L., Marchi, J. & Weitz, J. S. Metapopulation model of phage therapy of an acute Pseudomonas aeruginosa lung infection. mSystems 9, e0017124 (2024).

25. Marongiu, L., Burkard, M., Lauer, U. M., Hoelzle, L. E. & Venturelli, S. Reassessment of Historical Clinical Trials Supports the Effectiveness of Phage Therapy. Clin Microbiol Rev 35, e0006222 (2022).

26. GBD 2021 Antimicrobial Resistance Collaborators. Global burden of bacterial antimicrobial resistance 1990-2021: a systematic analysis with forecasts to 2050. Lancet 404, 1199–1226 (2024).

27. Chin, W. H. et al. Bacteriophages evolve enhanced persistence to a mucosal surface. Proc Natl Acad Sci U S A 119, e2116197119 (2022).

28. Joiner, K. L., Baljon, A., Barr, J., Rohwer, F. & Luque, A. Impact of bacteria motility in the encounter rates with bacteriophage in mucus. Sci Rep 9, 16427 (2019).

29. Sundberg, L.-R., Rantanen, N. & de Freitas Almeida, G. M. Mucosal Environment Induces Phage Susceptibility in Streptococcus mutans. Phage (New Rochelle) 3, 128–135 (2022).

30. Coelho, L. F. L. et al. Mucosal-adapted bacteriophages as a preventive strategy for a lethal Pseudomonas aeruginosa challenge in mice. Commun Biol 8, 13 (2025).

31. Almeida, G. M. F., Laanto, E., Ashrafi, R. & Sundberg, L.-R. Bacteriophage Adherence to Mucus Mediates Preventive Protection against Pathogenic Bacteria. mBio 10, (2019).

32. Almeida, G. M. de F. et al. Relevance of the bacteriophage adherence to mucus model for Pseudomonas aeruginosa phages. Microbiol Spectr 12, e0352023 (2024).

33. Wu, J. et al. Bacteriophage defends murine gut from Escherichia coli invasion via mucosal adherence. Nat Commun 15, 4764 (2024).

34. Barr, J. J. et al. Bacteriophage adhering to mucus provide a non-host-derived immunity. Proc Natl Acad Sci U S A 110, 10771–6 (2013).

35. Fu, K. et al. Escherichia coli phage ΦPNJ-9 adheres to mucus via a variant Hoc protein. J Virol 99, e0178924 (2025).

36. Pacios, O. et al. Molecular studies of phages-Klebsiella pneumoniae in mucoid environment: innovative use of mucolytic agents prior to the administration of lytic phages. Front Microbiol 14, 1286046 (2023).

37. Green, S. I. et al. Targeting of Mammalian Glycans Enhances Phage Predation in the Gastrointestinal Tract. mBio 12, (2021).

38. Sui, B. et al. Synergistic action of mucoactive drugs and phages against Pseudomonas aeruginosa and Klebsiella pneumoniae. Microbiol Spectr 13, e0160124 (2025).

39. de Freitas Almeida, G. M., Hoikkala, V., Ravantti, J., Rantanen, N. & Sundberg, L.-R. Mucin induces CRISPR-Cas defense in an opportunistic pathogen. Nat Commun 13, 3653 (2022).

40. Jakin Lazar, J. et al. Distinct effects of mucin on phage-host interactions in model systems of beneficial and pathogenic bacteria. Arch Virol 170, 133 (2025).

41. León, M. & Bastías, R. Virulence reduction in bacteriophage resistant bacteria. Front Microbiol 06, (2015).

42. Labrie, S. J., Samson, J. E. & Moineau, S. Bacteriophage resistance mechanisms. Nat Rev Microbiol 8, 317–27 (2010).

43. Castledine, M. et al. Parallel evolution of Pseudomonas aeruginosa phage resistance and virulence loss in response to phage treatment in vivo and in vitro. Elife 11, (2022).

44. Jung, H. C. et al. A distinct array of proinflammatory cytokines is expressed in human colon epithelial cells in response to bacterial invasion. J Clin Invest 95, 55–65 (1995).

45. Nagasaki, T. et al. Interleukin-6 released by colon cancer-associated fibroblasts is critical for tumour angiogenesis: anti-interleukin-6 receptor antibody suppressed angiogenesis and inhibited tumour-stroma interaction. Br J Cancer 110, 469–78 (2014).

46. Coutinho, H. D. M., Falcão-Silva, V. S. & Gonçalves, G. F. Pulmonary bacterial pathogens in cystic fibrosis patients and antibiotic therapy: a tool for the health workers. Int Arch Med 1, 24 (2008).

47. Lillehoj, E. P., Kim, B. T. & Kim, K. C. Identification of Pseudomonas aeruginosa flagellin as an adhesin for Muc1 mucin. American Journal of Physiology-Lung Cellular and Molecular Physiology 282, L751–L756 (2002).

48. Lillehoj, E. P. et al. Muc1 mucins on the cell surface are adhesion sites for Pseudomonas aeruginosa. American Journal of Physiology-Lung Cellular and Molecular Physiology 280, L181–L187 (2001).

49. Bucchioni, E., Kharitonov, S. A., Allegra, L. & Barnes, P. J. High levels of interleukin-6 in the exhaled breath condensate of patients with COPD. Respir Med 97, 1299–1302 (2003).

50. Reynolds, C. J. et al. Lung Defense through IL-8 Carries a Cost of Chronic Lung Remodeling and Impaired Function. Am J Respir Cell Mol Biol 59, 557–571 (2018).

51. Flynn, J. M., Niccum, D., Dunitz, J. M. & Hunter, R. C. Evidence and Role for Bacterial Mucin Degradation in Cystic Fibrosis Airway Disease. PLoS Pathog 12, e1005846 (2016).

52. Bonser, L. R. & Erle, D. J. Airway Mucus and Asthma: The Role of MUC5AC and MUC5B. J Clin Med 6, (2017).

53. Liyanage, T. D. et al. Biological Activity of Porcine Gastric Mucin on Stress Resistance and Immunomodulation. Molecules 25, (2020).

54. Marczynski, M. et al. Structural Alterations of Mucins Are Associated with Losses in Functionality. Biomacromolecules 22, 1600–1613 (2021).

55. Co, J. Y. et al. Mucins trigger dispersal of Pseudomonas aeruginosa biofilms. NPJ Biofilms Microbiomes 4, 23 (2018).

56. Haley, C. L., Kruczek, C., Qaisar, U., Colmer-Hamood, J. A. & Hamood, A. N. Mucin inhibits Pseudomonas aeruginosa biofilm formation by significantly enhancing twitching motility. Can J Microbiol 60, 155–66 (2014).

57. Chan, B. K. et al. Personalized inhaled bacteriophage therapy for treatment of multidrug-resistant Pseudomonas aeruginosa in cystic fibrosis. Nat Med 31, 1494–1501 (2025).

58. Roebuck, K. A. Regulation of interleukin-8 gene expression. J Interferon Cytokine Res 19, 429–38 (1999).

59. Tanaka, T., Narazaki, M. & Kishimoto, T. IL-6 in inflammation, immunity, and disease. Cold Spring Harb Perspect Biol 6, a016295 (2014).

60. Markwitz, P. et al. Emerging Phage Resistance in Pseudomonas aeruginosa PAO1 Is Accompanied by an Enhanced Heterogeneity and Reduced Virulence. Viruses 13, (2021).

61. Li, N. et al. Characterization of Phage Resistance and Their Impacts on Bacterial Fitness in Pseudomonas aeruginosa. Microbiol Spectr 10, (2022).

62. Pourcel, C., Midoux, C., Vergnaud, G. & Latino, L. The Basis for Natural Multiresistance to Phage in Pseudomonas aeruginosa. Antibiotics (Basel) 9, (2020).

63. Price-Whelan, A., Dietrich, L. E. P. & Newman, D. K. Pyocyanin alters redox homeostasis and carbon flux through central metabolic pathways in Pseudomonas aeruginosa PA14. J Bacteriol 189, 6372–81 (2007).

64. Huang, J., Sonnleitner, E., Ren, B., Xu, Y. & Haas, D. Catabolite Repression Control of Pyocyanin Biosynthesis at an Intersection of Primary and Secondary Metabolism in Pseudomonas aeruginosa. Appl Environ Microbiol 78, 5016–5020 (2012).

65. Welsh, M. A. & Blackwell, H. E. Chemical Genetics Reveals Environment-Specific Roles for Quorum Sensing Circuits in Pseudomonas aeruginosa. Cell Chem Biol 23, 361–369 (2016).

66. Lillehoj, E. P., Kato, K., Lu, W. & Kim, K. C. Cellular and molecular biology of airway mucins. Int Rev Cell Mol Biol 303, 139–202 (2013).

67. Freschi, L. et al. Clinical utilization of genomics data produced by the international Pseudomonas aeruginosa consortium. Front Microbiol 6, 1036 (2015).

68. Wiehlmann, L. et al. Population structure of Pseudomonas aeruginosa. Proc Natl Acad Sci U S A 104, 8101–6 (2007).

69. Cady, K. C., Bondy-Denomy, J., Heussler, G. E., Davidson, A. R. & O’Toole, G. A. The CRISPR/Cas adaptive immune system of Pseudomonas aeruginosa mediates resistance to naturally occurring and engineered phages. J Bacteriol 194, 5728–38 (2012).

