## Supplementary Figures for "Mucin-enriched environment shapes bacteriophage protection against *Pseudomonas aeruginosa* in airway epithelial cells"

Patricio, et al.

This file contains Supplementary Figures 1-3.

**Supplementary Figures**

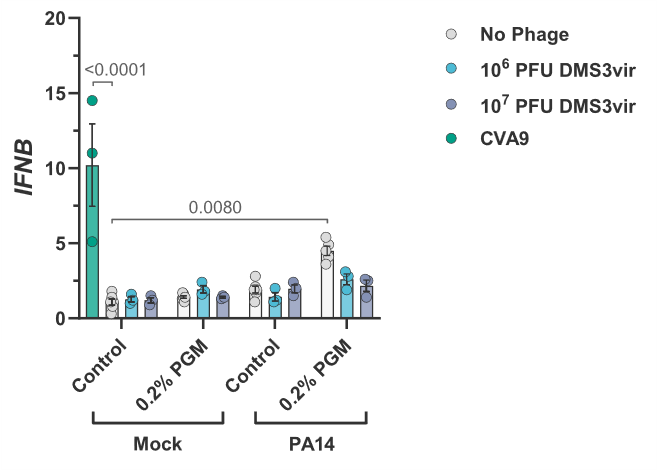

**Figure S1 | Phage DMS3vir does not induce type I interferon expression in P. aeruginosa-infected epithelial cells.** Relative expression of *IFNB* mRNA of A549 cells pre-incubated with DMS3vir and porcine gastric mucin (PGM) for 18 h, followed by infection with *P. aeruginosa* PA14 (1 x 10^5^ CFU) for 4 h. Quantitative data represent mean ± s.e.m. from at least two independent experiments. Coxsackievirus A9 (CVA9) infection (MOI 0.001) was used as a positive control. Statistical comparisons were performed using two-way ANOVA with multiple comparisons and Sidak’s post hoc test; p-values and replicate numbers are indicated.

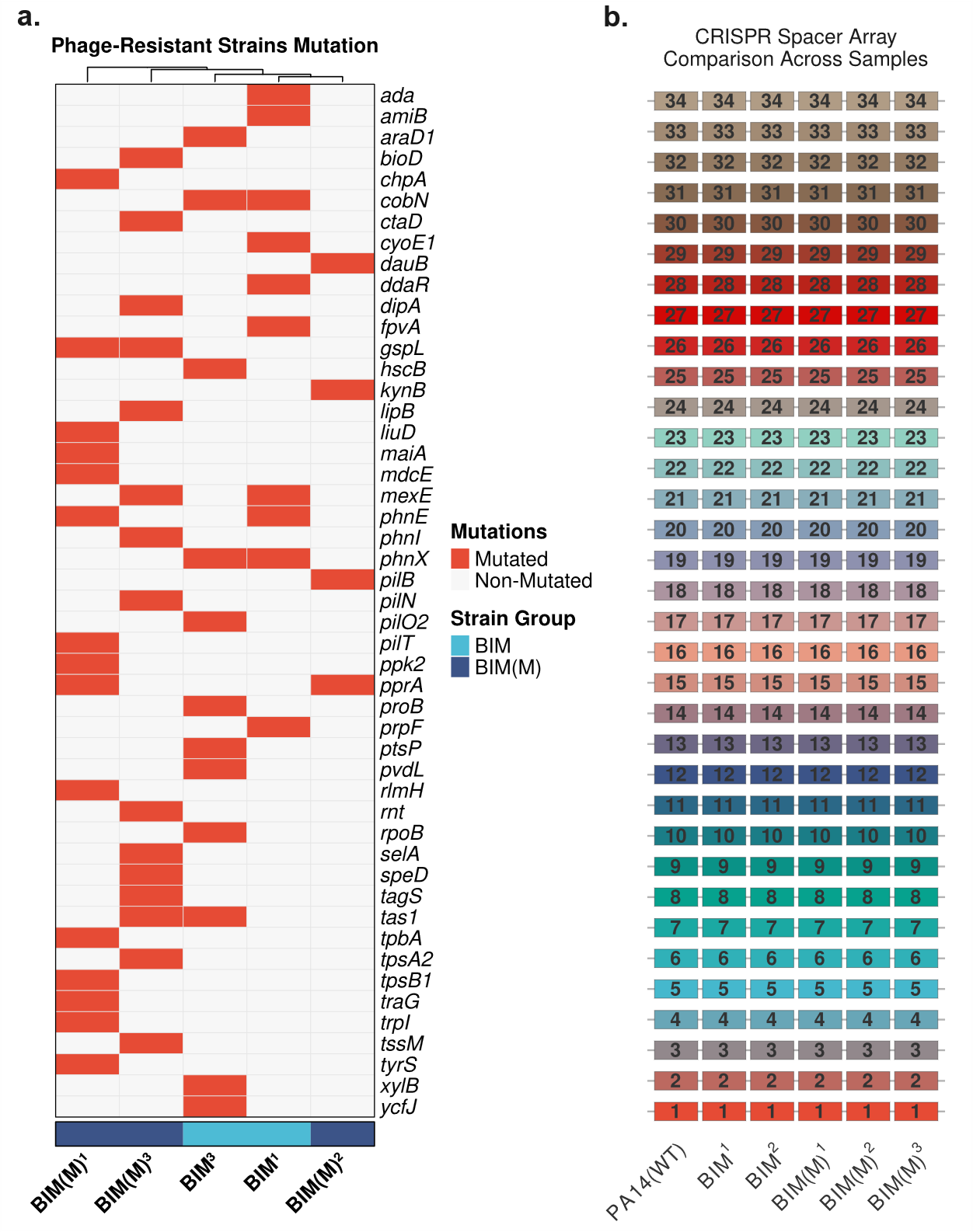

**Figure S2 | Whole genome sequencing analysis of bacteriophage-insensitive mutants (BIM) P. aeruginosa strains. (a)** Heatmap of mutated genes BIM and BIM(M) clones. Columns represent biological replicates. BIM and BIM(M) refers to phage-resistant clones isolated in the absence or presence of porcine gastric mucin (PGM), respectively, as described in Figure 7. Mutated genes shared with PA14(WT), PA14(C), or PA14(M) were excluded. **(b)** CRISPR spacer acquisition analysis.

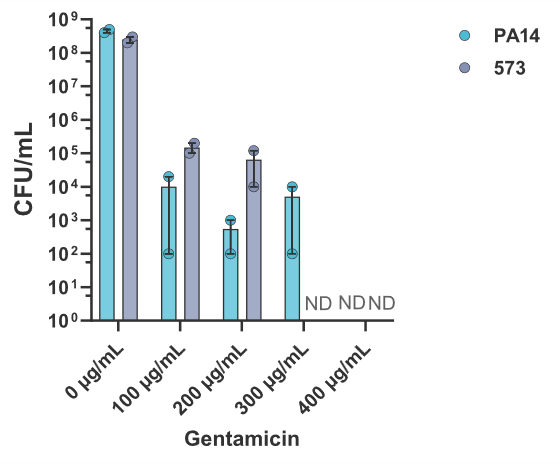

**Figure S3 | Antibacterial effects of gentamicin on *P. aeruginosa*.** *P. aeruginosa* strains PA14 and 573 exposed to gentamicin for 4 h. Antimicrobial effects of gentamicin were analyzed by bacterial colony-forming unit (CFU) quantification. ND: Non-detected. Statistical comparisons were performed using two-way ANOVA with multiple comparisons and Sidak’s post hoc test; p-values and replicate numbers are indicated.
