## Supplementary material for "Mucin-enriched environment shapes bacteriophage protection against *Pseudomonas aeruginosa* in airway epithelial cells": Additional Method Description on Bioinformatic Analysis.docx

**Patricio, et al.**

1. **RNA Sequencing and Analysis**
   1.1. **Sample Preparation and Sequencing**

Total RNA was extracted from A549 human lung carcinoma cells (n=3 biological replicates per group: untreated control and PGM-treated) using the RNeasy Mini Kit (Qiagen) according to the manufacturer's protocol. RNA integrity and quality were assessed using the Agilent 2100 Bioanalyzer, with all samples exhibiting an RNA Integrity Number (RIN) > 8. Strand-specific mRNA libraries were prepared from poly(A)-enriched RNA and sequenced on the DNBSEQ-G400 platform (BGI Tech) using paired-end 150 bp reads (PE150), targeting ~20–30 million reads per sample. Raw FASTQ files were deposited in the NCBI Sequence Read Archive (SRA) under BioProject accession PRJNA1308451.

1.2. **Data Processing and Quality Control**

Upstream processing followed the Chipster RNA-seq workflow (version accessed October 2025; <https://chipster.csc.fi/manual/rna-seq-summary.html>). Adapter trimming and quality filtering were done beforehand by BGI Tech (see Supplementary Information). Raw reads quality control was confirmed using FastQC (v0.12.1) to evaluate per-base sequence quality, adapter content, and overrepresented sequences. Sequencing summaries were generated with MultiQC (v1.31). Reads were aligned to the human reference genome Homo_sapiens.GRCh38 using STAR (v2.7.11b) with default parameters. Alignment quality was evaluated using Picard CollectMultipleMetrics (v3.2.0) for metrics including alignment rate, junction saturation, and 5'/3' bias. Gene-level read counts were quantified using HTSeq (v2.0.3) with default parameters against the GRCh38 GTF annotation, producing per-sample TSV files. These counts were combined into a matrix and metadata file using tidyverse (v2.0.0) in R (v4.4.1). MD5 checksums were generated for all input count files. Count-aligned read files were merged into one count table using the Chipster tool ‘Define NGS experiment’. Differential expression (DE) analysis was performed using both DESeq2 (v1.48.2) and edgeR (v4.6.3) with default parameters. Genes significant in both tools (FDR < 0.05) were intersect via Venn diagram using ‘identifier’ as common denominator.

1.3. **Gene Annotation**

Using R, significant genes (Ensembl IDs) were annotated using biomaRt (v2.62.1) against the Ensembl human dataset to retrieve HGNC symbols. Annotations were merged with org.Hs.eg.db (v3.20.0) for Entrez ID mapping. Duplicate symbols were resolved by retaining the entry with the highest absolute log2FC. The average log2 fold changes (logFC) across replicates were classified as upregulated (logFC > 0.5) and downregulated (logFC < -0.5) genes.

1.4. **Gene Ontology Enrichment**

GO Biological Process (BP) enrichment (qvalueCutoff = 0.05) was performed separately for upregulated (logFC > 0.5) and downregulated (logFC < -0.5) gene sets using clusterProfiler (v4.14.6) with org.Hs.eg.db. Enriched terms were manually curated (e.g., filtering for relevance). The terms were manually grouped into categories (Stress & Inflammation, Metabolic Reprogramming, Adhesion/Junctions & Plasticity, Growth-Factor Signaling) for interpretive visualization and did not alter statistical results.

1.5. **Visualization**

All visualizations were generated in R using ggplot2 (tidyverse v2.0.0), ggrepel (v0.9.6), ggnewscale (v0.5.1), ggtext (v0.1.2), and DESeq2; tidyverse, ComplexHeatmap (v2.22.0) and circlize (v0.4.16) for heatmaps; dendextend (v1.19.0) for dendrogram compression. For Principal Component Analysis (PCA), variance-stabilizing transformation (VST) was applied to the count matrix using DESeq2. The top 1,000 most variable genes (by row variance) were selected. PC1 and PC2 (percent variance explained) were plotted by replicates, repel text, and annotated by group (Control; PGM). For volcano plot, annotated DE genes were plotted as -log10(FDR) vs logFC. Labels for top 10 |logFC| genes using ggrepel. For differential expression heatmap, Z-score normalized log2 fold changes (relative to control average) for top DE genes (|logFC| > 1) were clustered (Euclidean distance; Ward's method) with compressed dendrograms. Rows were split by regulation (Upregulated/Downregulated). For GO enrichment bar plot, curated GO terms were plotted as Fold Enrichment (x-axis) vs. Description (y-axis), with bars colored by -log10(FDR) (upregulated and downregulated). Tiles were annotated by functional group. Gene lists were displayed as textboxes. Terms were ordered by regulation and descending Fold Enrichment.

2. **Whole-Genome Sequencing and Analysis**
   2.1. **Sample Preparation and Sequencing**

Genomic DNA was extracted from Pseudomonas aeruginosa PA14 strains (PA14(WT), PA14(C), PA14(M), BIM^1^, BIM^2^, BIM^3^, BIM(M)^1^, BIM(M)^2^, BIM(M)^3^; n=1 per strain) using the DNeasy Blood & Tissue Kit (Qiagen, cat. 69504) per manufacturer's instructions. DNA concentration and purity were quantified using the Qubit dsDNA HS Assay on a Qubit 4 fluorometer (Invitrogen). Paired-end libraries (insert sizes ≤800 bp) were prepared and sequenced on the DNBSEQ-G400 platform (BGI Tech) using PE150 reads (~50–100x coverage target). Sequencing BIM^2^ failed due to technical issues and was excluded. Raw FASTQ files were deposited in the NCBI SRA under BioProject PRJNA1308451.

2.2. **Data Processing, Alignment, and Quality Control**

Reads were quality-filtered with fastp (v0.23.4) and summarized with MultiQC. Alignments to the PA14 reference were generated with BWA-MEM (v0.7.19) piped to SAMtools (v1.21) for sorting and indexing. Duplicates were marked and removed with GATK MarkDuplicates (v4.6.2). Read groups were added via GATK AddOrReplaceReadGroups. Base quality score recalibration was not applied. Pileup files were generated with SAMtools mpileup. Variants (SNPs and Indels) were called with VarScan (v2.4.2). SNP and indel VCF files were converted to TSVs with bcftools query (v1.12) and combined to a single file. Whole genome assemblies for CRISPR detection were generated de novo with SPAdes (v4.0.0).

2.3. **Variant Annotation and Filtering**

Variants were combined across samples into a long-format using tidyverse, with metadata for groups (Control: PA14(WT/C/M); BIM: BIM^1^/^3^; BIM(M): BIM(M^1^/M^2^/M^3^)). Wild-type (PA14_WT) positions were excluded via anti-join. Coding consequences were predicted using VariantAnnotation (v1.52.0) and GenomicRanges (v1.52.0): VRanges objects were created from variants, aligned to the PA14 GFF-derived TxDb (via txdbmaker v1.70.0 and rtracklayer v1.60.3), and annotated with predictCoding against the reference FASTA (Biostrings v2.68.0). Ambiguous reference bases were filtered. Overlaps with genes (GRanges from GFF) assigned GeneID and gene names were merged from GFF and UniProt annotation. Non-synonymous variants were retained and control-shared positions were further excluded. Variant types (SNP/Insertion/Deletion/Complex) were classified by REF/ALT lengths. MD5 checksums were generated for TSVs. Annotated gene’s function/pathway was manually curated at <https://www.kegg.jp/kegg/kegg2.html>, accessed September 2025.

2.4. **CRISPR Spacer Array Detection**

CRISPR arrays were detected in SPAdes contigs using minced (v0.2.0). Lines were regex-matched for positions, repeats, and spacers. Spacers were filtered (non-NA), and overlapping or redundant arrays were merged by locus coordinates. Canonical sequences were computed (forward, reverse complement) using Biostrings. Arrays were oriented relative to PA14(WT) (forward/reverse/unknown) and combined to a single file.

2.5. **Visualization**

Visualizations used ComplexHeatmap, circlize, dendextend, tidyverse, ggtext, and viridis (v0.6.5); ggsci (v3.2.0) for palettes. For binary mutation heatmap, non-synonymous variants were pivoted to a binary matrix (1 = mutated, 0 = non-mutated). Genes with NA or not-annotated (PA14_XXXX) GeneName were filtered. Hierarchical clustering was performed using Euclidean distance and Ward’s linkage. Strain groups (BIM and BIM(M) was annotated. For kegg pathway stacked bar plot, annotated genes were merged with kegg.tsv via GeneName. Counts per Classification/Sample were stacked and genes labeled. The CRISPR spacer acquisition tile plot was tiled by Sample vs. SpacerNum.
