## Supplementary material for "Mucin-enriched environment shapes bacteriophage protection against *Pseudomonas aeruginosa* in airway epithelial cells": Description of Additional Supplementary Files.docx

File Name: Supplementary Data 1

Description: Differential expression data of A549 cells exposed to porcine gastric mucin.

File Name: Supplementary Data 2

Description: GO enrichment data A549 cells exposed to porcine gastric mucin.

File Name: Supplementary Data 3

Description: Whole-genome sequencing data of *Pseudomonas aeruginosa* PA14 isolates mutations.

File Name: Supplementary Data 4

Description: Kegg function/pathway annotation for mutated genes in *Pseudomonas aeruginosa* PA14 isolates.

File Name: Supplementary Data 5

Description: CRISPR spacers acquisition data of *Pseudomonas aeruginosa* PA14 isolates.
